# A tumor-cell MHC-II program is associated with checkpoint-blockade outcomes across stages of bladder cancer

**DOI:** 10.64898/2026.09.21.753036

**Authors:** Joaquim Bellmunt, Yingtian Xie, Miguel Gomez Munoz, Shweta Kukreja, Sarah Walker, Ruiyang He, Rong Li, Xintao Qiu, Tao Zhang, Paloma Rodriguez, Diego García González, Ilana Epstein, Julia Parera, Nuria Juanpere, Marta Lorenzo, Oscar Buisan, Jordi Senserrich, Pol Servián, Tatiana Silva, Myles Brown, Cecilia Cabrera, Paloma Cejas, Henry W. Long

**Affiliations:** Harvard Medical School, Boston, MA, USA; Lank Center for Genitourinary Oncology, Dana-Farber Cancer Institute, Boston, MA, USA; Center for Functional Cancer Epigenetics, Dana-Farber Cancer Institute, Boston, MA 02215, USA; Department of Pathology, Hospital del Mar, Barcelona, Spain; Department of Urology, IDIBELL Hospital Universitario de Bellvitge, Barcelona, Spain; Irsicaixa, Hospital Universitari Germans Trias i Pujol, Badalona, Barcelona, Spain; Department of Urology, Hospital Universitari Germans Trias i Pujol, Badalona, Spain; Germans Trias I Pujol Research Institute (IGTP), Badalona, Spain; Autonomous University of Barcelona, Bellaterra, Barcelona, Spain; Translational Oncology Laboratory, Hospital La Paz Institute for Health Research (IdiPAZ), Madrid, Spain

## Abstract

**Purpose:** A minority of patients with bladder cancer derive lasting benefit from immunotherapy, and current biomarkers capture only part of the biology. We evaluated whether tumor-cell MHC class II defines an immune state associated with outcome across disease stages and after PD-1/PD-L1 blockade.

**Experimental Design:** We localized differential MHC-II activity to malignant epithelial cells by single-nucleus and single-cell RNA sequencing and confirmed HLA-DR expression by immunohistochemistry in 122 specimens. Chromatin profiling and IFN-γ stimulation of cell lines and primary tumors supported derivation of an 11-gene tumor-cell program. Associations were tested in 434 patients with non-muscle-invasive disease, 82 receiving neoadjuvant pembrolizumab, and 288 receiving atezolizumab for metastatic disease.

**Results:** Tumor-cell MHC-II was present in approximately one third of bladder cancers, and malignant cells accounted for the differential signal between MHC-II-high and MHC-II-low tumors. The program was inducible by IFN-γ through JAK/STAT signaling. Within luminal non-muscle-invasive disease, MHC-II-low status was associated with progression (HR 3.31, 95% CI 1.37 to 7.99). In PURE-01, the 11-gene program was associated with pathological complete response (52% versus 24%, p = 0.018) and recurrence-free survival (p = 0.0054), retaining an independent association in models including tumor mutational burden and PD-L1. In bladder-primary metastatic disease, program-high status was associated with overall survival in unadjusted analysis (HR 0.62, 95% CI 0.44 to 0.89); the association did not extend to upper tract tumors (interaction p = 0.0018).

**Conclusions:** Tumor-cell MHC-II defines an inducible state associated with outcomes across bladder cancer stages and after checkpoint blockade, supporting prospective evaluation by RNA and immunohistochemistry.

**Translational Relevance:** Bladder cancer is one of few malignancies in which immunotherapy is given from non-muscle-invasive through metastatic disease, yet tissue-based biomarkers identify only a subset of patients who derive benefit. We found approximately one third of bladder tumors express MHC class II on malignant epithelial cells. Within luminal non-muscle-invasive disease, MHC-II-low status marked a threefold higher progression risk. In checkpoint-treated muscle-invasive and bladder-primary metastatic disease, an 11-gene tumor-cell program was associated with response and survival; in the neoadjuvant cohort, it added information beyond tumor mutational burden and PD-L1. Complementary assays, amenable to clinical-laboratory implementation, capture different aspects of this state: bulk RNA was used in the outcome analyses, whereas HLA-DR immunohistochemistry demonstrated tumor-cell protein expression in a 122 specimen TMA. Because the program is inducible, a negative result may identify tumors lacking inflammatory input rather than tumors incapable of responding. As enfortumab vedotin plus pembrolizumab becomes standard of care, tsMHC-II is a candidate biomarker of the immunotherapy-sensitive component of such regimens and warrants prospective evaluation across the treatment course.

## INTRODUCTION

Over the past decade, immune checkpoint blockade, particularly targeting the PD-1/PD-L1 axis, has transformed the management of multiple solid tumors, including melanoma, lung cancer, and renal cell carcinoma(1–3). Bladder cancer is one of the few malignancies treated with immune-directed therapies across the disease course.

Intravesical Bacillus Calmette-Guérin (BCG) remains standard of care for high-risk non-muscle-invasive bladder cancer (NMIBC), which accounts for approximately 70% of new diagnoses, and acts in part by engaging CD4⁺ T-cell responses(4–6), whereas PD-1/PD-L1 inhibitors have produced durable responses in advanced urothelial carcinoma(7–9).

The treatment landscape has shifted toward combinations, with enfortumab vedotin plus pembrolizumab now first-line standard of care in advanced disease(10) and checkpoint inhibitor monotherapy reserved for selected patients. Defining the tumor-intrinsic state associated with checkpoint blockade sensitivity therefore remains relevant both where immunotherapy is given alone, including the adjuvant and non-muscle-invasive settings, and as the immunotherapy component of combination regimens as only a minority of patients achieve lasting benefit from either modality, and PD-L1 immunohistochemistry and tumor mutational burden capture only part of the underlying biology. A clinically useful biomarker would therefore need to identify the immune-responsive tumor state across settings in which treatment and clinical endpoints differ.

One candidate is tumor-intrinsic expression of MHC class II (MHC-II). Although MHC-II expression is typically restricted to professional antigen-presenting cells, multiple tumor types can express MHC-II under inflammatory conditions, primarily through interferon-γ (IFN-γ)-mediated induction of the class II transactivator CIITA(11–13), with chromatin accessibility at MHC-II loci as a key permissive factor. Tumor-specific MHC-II expression (tsMHC-II) enables direct presentation of tumor antigens to CD4⁺ T cells, supporting helper T-cell responses, CD8⁺ T-cell priming and persistence, and in some contexts direct CD4⁺ cytotoxic activity against tumor cells(14–16). In melanoma, tsMHC-II is a tumor-autonomous phenotype that predicts response to PD-1/PD-L1 blockade but not to CTLA-4 inhibition(17,18). In bladder cancer, Yi et al.(19) defined a bulk transcriptional signature associated with T-cell infiltration, IFN-γ signaling, and improved survival after PD-L1 blockade, supporting the clinical relevance of this pathway while leaving its cellular source unresolved.

Whether tsMHC-II can support a clinically useful biomarker in bladder cancer depends on three translational requirements. First, biological specificity: the signal must be localized to malignant cells rather than infiltrating antigen-presenting cells. Second, clinical association: the tumor-cell state should be tested against outcomes in independent checkpoint-treated settings rather than inferred from immune correlation alone. Third, assay feasibility: the RNA-defined state should be detectable in routine tissue using a practicable readout that retains tumor-cell localization. The prevalence and clinical relevance of tsMHC-II across bladder cancer stages, particularly in NMIBC, remain poorly defined. A recent single-cell study of muscle-invasive disease interpreted tumor MHC-II as supporting CD8⁺ T-cell exhaustion and immune evasion(20), leaving unresolved whether the phenotype marks immune-active or immune-suppressed disease.

Here we evaluate tsMHC-II against these translational requirements. We first show that approximately one third of bladder tumors, spanning NMIBC and muscle-invasive disease, express MHC-II directly on malignant epithelial cells, and that these cells account for the differential pathway signal in bulk RNA-seq, establishing tumor-cell specificity. We show that tsMHC-II resolves prognostic heterogeneity within luminal NMIBC, and an 11-gene tumor-cell program is associated with response and survival after checkpoint blockade in independent neoadjuvant and metastatic settings. In the neoadjuvant cohort, the association persists in models incorporating tumor mutational burden and PD-L1, while in the metastatic setting its lack of association in upper tract tumors suggests an anatomic boundary. HLA-DR immunohistochemistry demonstrates tumor-cell protein expression in tissue, complementing the bulk RNA score used in the outcome analyses. We further show that the program is inducible and IFN-γ-dependent, licensed by chromatin that is already accessible in most bladder tumors, indicating that a negative test may reflect absent inflammatory input rather than fixed incapacity. A discrete IFN-γ-producing, heat-shock-enriched CD8-lineage T-cell subset provides a potential intratumoral source of IFN-γ. Together, these findings support prospective evaluation of tsMHC-II for bladder cancer immunotherapy stratification.

## RESULTS

### Tumor-cell MHC-II is a prevalent, protein-detectable feature of bladder cancer at both early and advanced stages

#### MHC-II pathway activity is bimodally distributed in bladder cancer across disease stages

Our recent chromatin profiling of NMIBC clinical samples identified striking H3K27ac enrichment at the MHC-II gene cluster on chromosome 6 in luminal tumors, with promoter peak intensities comparable to those of established tumor-expressed reference genes(21) (Supplementary Figure 1a). To ask whether this active-chromatin signal corresponds to a definable expression state, we examined four bladder cancer cohorts spanning early- and late-stage disease, showing the largest cohort of each stage in the main figure and the remaining two in the supplement. Using Gene Set Variation Analysis (GSVA)(22), we computed an MHC-II pathway score per sample in each cohort.

In both NMIBC cohorts, GSVA enrichment scores displayed a distinct bimodal distribution, mirroring patterns previously described in melanoma(17). This bimodality was evident in the large UROMOL cohort (Figure 1a, left) and reproduced in a previously published internal dataset (the Bellmunt cohort; Supplementary Figure 1b, right). We modeled each distribution as a two-component Gaussian mixture and used the inter-peak minimum as a data-driven cutpoint to assign MHC-II–high versus MHC-II–low status (Methods). In both cohorts, MHC-II–high tumors were significantly enriched for the luminal (Class 2a) subtype as defined by the UROMOL2021 classifier(23) (UROMOL: p = 8.6×10⁻⁸; Bellmunt: p = 1.6×10⁻⁴; Fisher’s exact test).

**Figure 1.**
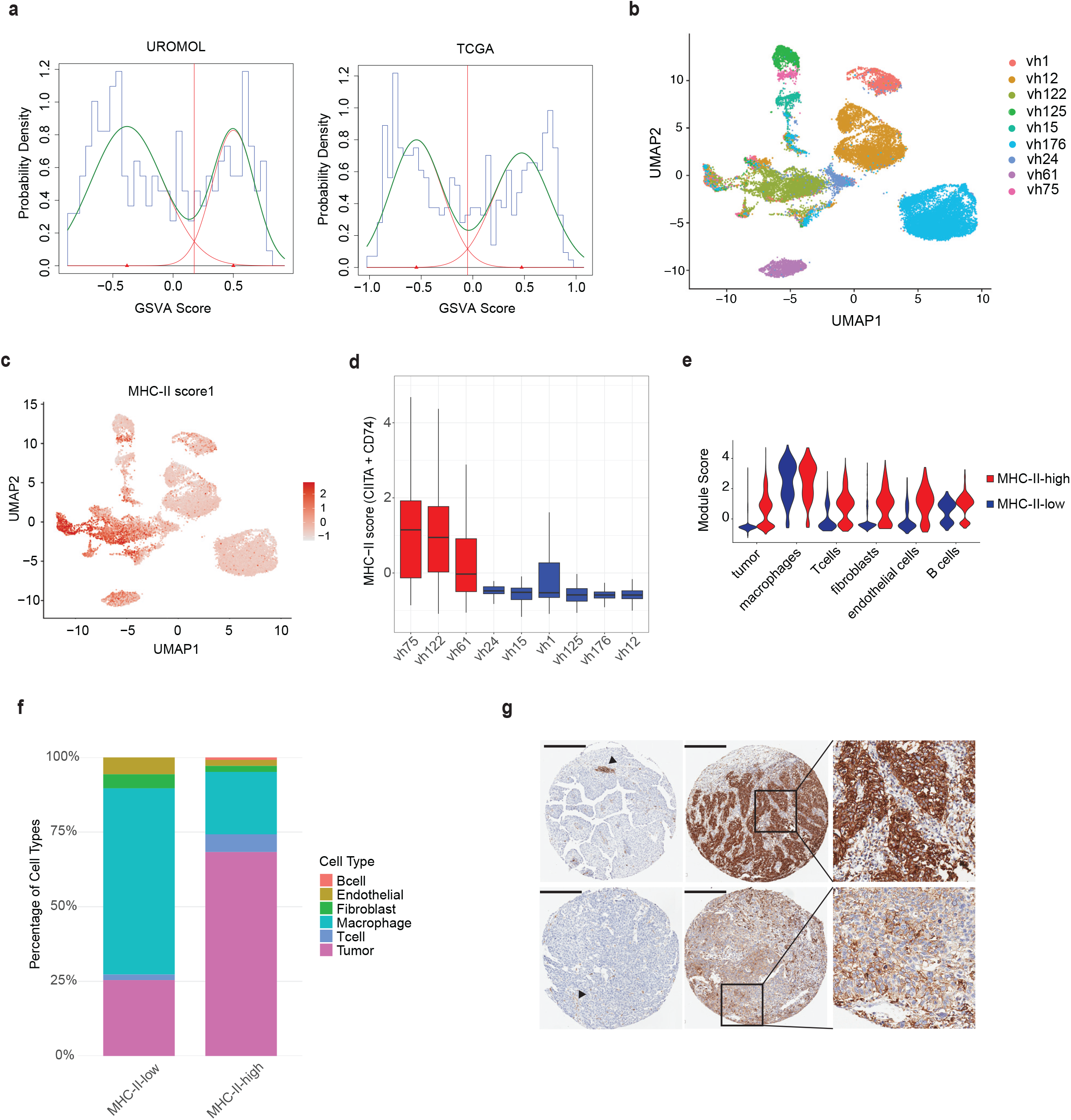
Tumor-intrinsic MHC-II pathway activation is prevalent across bladder cancer disease stages. **(a)** Distribution of GSVA-derived MHC-II pathway signature scores in the UROMOL NMIBC cohort (n = 475; left) and the TCGA MIBC cohort (n = 410; right). Histograms display the distribution of enrichment scores across all samples. Green curves indicate fitted bimodal Gaussian mixture models; the red vertical line marks the inter-peak minimum used as a data-driven cutpoint to classify tumors as MHC-II–high or MHC-II– low (see Methods). The bimodal distribution indicates that MHC-II pathway activity segregates bladder cancers into two discrete transcriptional states. **(b)** UMAP visualization of 23,136 nuclei from the internal snRNA-seq cohort (nine NMIBC), showing only cells annotated as malignant epithelial cells. Points are colored by patient identity (n = 9 samples). Cell type annotation was performed using canonical marker genes and inferCNV-based copy-number variation analysis (see Methods). **(c)** UMAP visualization of tumor cells in the internal snRNA-seq cohort, colored by combined *CD74* and *CIITA* expression (summed normalized counts). High co-expression identifies a subset of malignant cells with active MHC-II transcription, corresponding to patients classified as MHC-II–high. **(d)** Box plot showing per-sample MHC-II expression scores (*CIITA* + *CD74*, summed across tumor cells) for each of the nine NMIBC samples in the internal cohort. Samples classified as MHC-II–high are shown in red; MHC-II–low samples are shown in blue. Classification was based on bimodal thresholding of GSVA scores in matched bulk RNA-seq data. Three of nine samples (33%) were classified as MHC-II–high, matching the tumor-specific prevalence measured by HLA-DR immunohistochemistry in the independent TMA cohort (31%, panel g). **(e)** Violin plots showing expression of MHC-II module score across annotated cell types in the internal snRNA-seq cohort split by MHC-II high/low. Each violin represents the distribution of normalized gene expression values within a given cell type. MHC-II pathway components are highly expressed in macrophages and B cells, as expected for professional antigen-presenting cells, but are also detectable in a subset of malignant epithelial cells. **(f)** Proportional cell-type composition of MHC-II–positive cells in MHC-II–low versus MHC-II–high tumors in the internal snRNA-seq cohort. The fraction of MHC-II–positive cells of tumor origin rises from 26% in MHC-II–low tumors to 68% in MHC-II–high tumors, with a reciprocal decrease in the macrophage fraction. **(g)** Four representative immunohistochemistry (IHC) images of MHC-II staining from a bladder cancer tumor microarray (TMA; n = 122 evaluable specimens). The scale bars are 300 microns each. Left: MHC-II–negative cases showing no staining in tumor cells. Arrows point to staining in immune cells. Right: MHC-II–positive cases including expanded views show extensive staining of tumor cells. Staining was performed with a validated DAKO HLA-DR antibody and scored by an expert genitourinary pathologist using a histoscore-based method (see Methods).

The same pattern was evident in MIBC. Both the TCGA cohort (Figure 1a, right) and the PURE-01 cohort (Supplementary Figure 1b, left) displayed bimodal MHC-II GSVA score distributions. Together, these analyses identify two discrete transcriptional states of MHC-II pathway activity that recur across non-muscle-invasive and muscle-invasive disease, one with active and one with inactive MHC-II transcription. Because this bulk GSVA score aggregates signals from every cell in the tumor specimen, we next used single-cell profiling to determine which cells give rise to it.

#### Malignant epithelial cells, not immune infiltrate, account for the differential MHC-II signal

Bulk transcriptomic profiling cannot distinguish whether MHC-II pathway activity originates from tumor cells or from infiltrating immune cells, both of which can express MHC-II in inflamed microenvironments. To resolve the cellular source of the signal, we performed single-nucleus RNA sequencing (snRNA-seq) on nine NMIBC samples from our previous study(21). Samples were profiled from FFPE tissue using 10x Genomics FLEX chemistry, and cell types were assigned by canonical marker expression and inferCNV-based copy-number profiling (Methods). After quality filtering, 26,851 nuclei across the nine samples were retained for downstream analysis of which 23,136 were tumor nuclei (Figure 1b). The FLEX probe panel provides reliable coverage of *CD74* and *CIITA* but limited coverage of the broader MHC-II gene set, so we used the summed expression of *CD74* and *CIITA* as a proxy for pathway activity in this cohort; this proxy correlates strongly with the full pathway score (Pearson r = 0.90, p < 0.001) in an external 3′ scRNA-seq dataset described below (Supplementary Figure 1e).

As expected, MHC-II pathway activity was readily detectable in professional antigen-presenting cells, with the highest per-cell scores in macrophages and B cells (Figure 1e). Within the malignant epithelial compartment, clear pathway activation was present in three of the nine NMIBC samples (Figure 1c, d), a proportion consistent with the prevalence observed in the bulk cohorts. All three MHC-II–high samples were classified as luminal-like using the in-house NMIBC molecular framework derived from the same cohort(21), consistent with the luminal enrichment observed in the bulk classification.

To determine which cells account for the difference in MHC-II pathway activity between MHC-II–high and MHC-II–low tumors, we decomposed the per-cell MHC-II score by compartment. Immune and stromal cells expressed MHC-II at the highest per-cell levels in both groups, increasing further between MHC-II–low and MHC-II–high tumors (mean 0.59 versus 1.43; Supplementary Figure 1g). Tumor cells, by contrast, underwent a qualitative shift, with their mean score moving from negative in MHC-II–low tumors to clearly positive in MHC-II–high tumors (mean −0.52 versus 0.62). Because tumor cells vastly outnumber immune and stromal cells in these specimens, this shift converted the tumor compartment from a net-negative to a net-positive contributor and dominated the net change in total MHC-II signal between the two groups (Supplementary Figure 1h). Consistent with this, the proportion of MHC-II–positive cells of tumor origin rose from 26% in MHC-II–low tumors to 68% in MHC-II–high tumors, with a reciprocal decline in the macrophage fraction (Figure 1f).

Tumor cells are therefore the principal source of the *differential* MHC-II signal that distinguishes MHC-II–high from MHC-II–low bladder cancers, even though professional antigen-presenting cells retain higher per-cell expression. The bimodal MHC-II signal detected in bulk RNA-seq reflects this switch in tumor-cell expression rather than variation in immune-infiltrate composition.

To test whether this finding extends to muscle-invasive disease in a larger cohort, we analyzed a published MIBC scRNA-seq dataset of 25 cases profiled with 10x Genomics 3′ chemistry(24), processed through the same analytical pipeline (Methods). After quality filtering, 67,988 cells were retained (Supplementary Figure 1c). 6 of 25 patients showed extensive tumor-intrinsic MHC-II expression (Supplementary Figure 1d, f), consistent with the prevalence observed in the NMIBC cohort. Together, these single-cell data identify malignant epithelial cells as the dominant source of the differential MHC-II signal in bladder tumors across both disease stages.

#### Immunohistochemistry detects tumor-cell MHC-II protein in one third of bladder cancers

Transcriptomic and single-cell evidence cannot directly resolve whether the MHC-II protein complex is expressed on the tumor-cell surface, where it would be available for T-cell engagement. To confirm tumor-cell protein expression in an independent patient cohort, we performed immunohistochemistry (IHC) for MHC-II on a tumor microarray (TMA) comprising 122 evaluable bladder cancer specimens(25). Sections were stained with an HLA-DR antibody (clone CR3/43) previously validated for tumor MHC-II detection in melanoma(18) and scored by an expert genitourinary pathologist blinded to clinical and molecular annotations using a histoscore-based method (Figure 1g; Methods).

Tumor-cell membrane MHC-II expression (histoscore > 2) was detected in 31% of evaluable cases overall (38/122), with comparable rates in NMIBC (32/107, 30%) and the smaller MIBC subset (6/15, 40%). This protein-level prevalence closely matches the tumor-specific frequency identified by snRNA-seq (33% of internal NMIBC samples), confirming at the protein level that approximately one-third of bladder cancers express tumor-cell MHC-II.

The bulk and tumor-specific measures do not report the same quantity, and the distinction determines how prevalence should be stated. Tumor-specific measures agree closely: 31% by HLA-DR immunohistochemistry, 33% by snRNA-seq in the internal NMIBC cohort, and 24% (6 of 25) in the external MIBC single-cell cohort. Bulk GSVA classification assigns a higher proportion of tumors to the MHC-II-high group in every cohort (UROMOL 47.7%, Bellmunt 50%, TCGA 50.1%, PURE-01 40.2%, and 45.8% in IMvigor210). The difference is expected. Bulk scores aggregate MHC-II expression from every cell in the specimen, including professional antigen-presenting cells, which express MHC-II constitutively and at higher per-cell levels than tumor cells, so in tumors where malignant cells are MHC-II-negative the immune compartment alone can place a specimen near or above the mixture cutpoint. Consistent with this, only 26% of MHC-II-positive cells are of tumor origin in MHC-II-low tumors, compared with 68% in MHC-II-high tumors (Figure 1f). Bulk classification therefore identifies tumors in which the pathway is active somewhere in the specimen, while immunohistochemistry and single-cell profiling identify tumors in which malignant cells themselves express the protein. We use the tumor-specific figure of approximately one third as the prevalence of tsMHC-II throughout, and the bulk-derived signature retains clinical association because the tumor-cell contribution dominates the differential signal between groups.

Together, bulk transcriptomics established the bimodal MHC-II signal across four independent cohorts, while single-cell profiling and IHC localize it to malignant epithelium and establish tumor-cell MHC-II as a prevalent feature of both early- and late-stage bladder cancer.

### Tumor-cell MHC-II is an inducible state licensed by IFN-γ rather than a fixed tumor property

Prior work in melanoma and breast cancer has shown that IFN-γ-driven MHC-II induction depends on chromatin accessibility at CIITA and MHC-II promoters, and that epigenetic repression can uncouple inflammatory signaling from transcriptional activation(18,26–28). To test this regulatory architecture in bladder cancer, we profiled basal, luminal, and mixed-subtype cell lines (Supplementary Table 1) at baseline and after IFN-γ stimulation, generating RNA-seq for six lines and ATAC-seq for five.

#### MHC-II loci are accessible but transcriptionally silent in most bladder tumors and cell lines

At baseline, ATAC-seq showed that most cell lines had accessible chromatin at MHC-II regulatory loci, including CD74, CIITA, HLA-DRA, HLA-DRB1, HLA-DQB1, and HLA-DPA1 (Figure 2a). This permissive state occurred across molecular subtypes, suggesting that these loci are open without inflammatory stimulation. J82 and SW-780 were exceptions, with minimal baseline accessibility.

**Figure 2.**
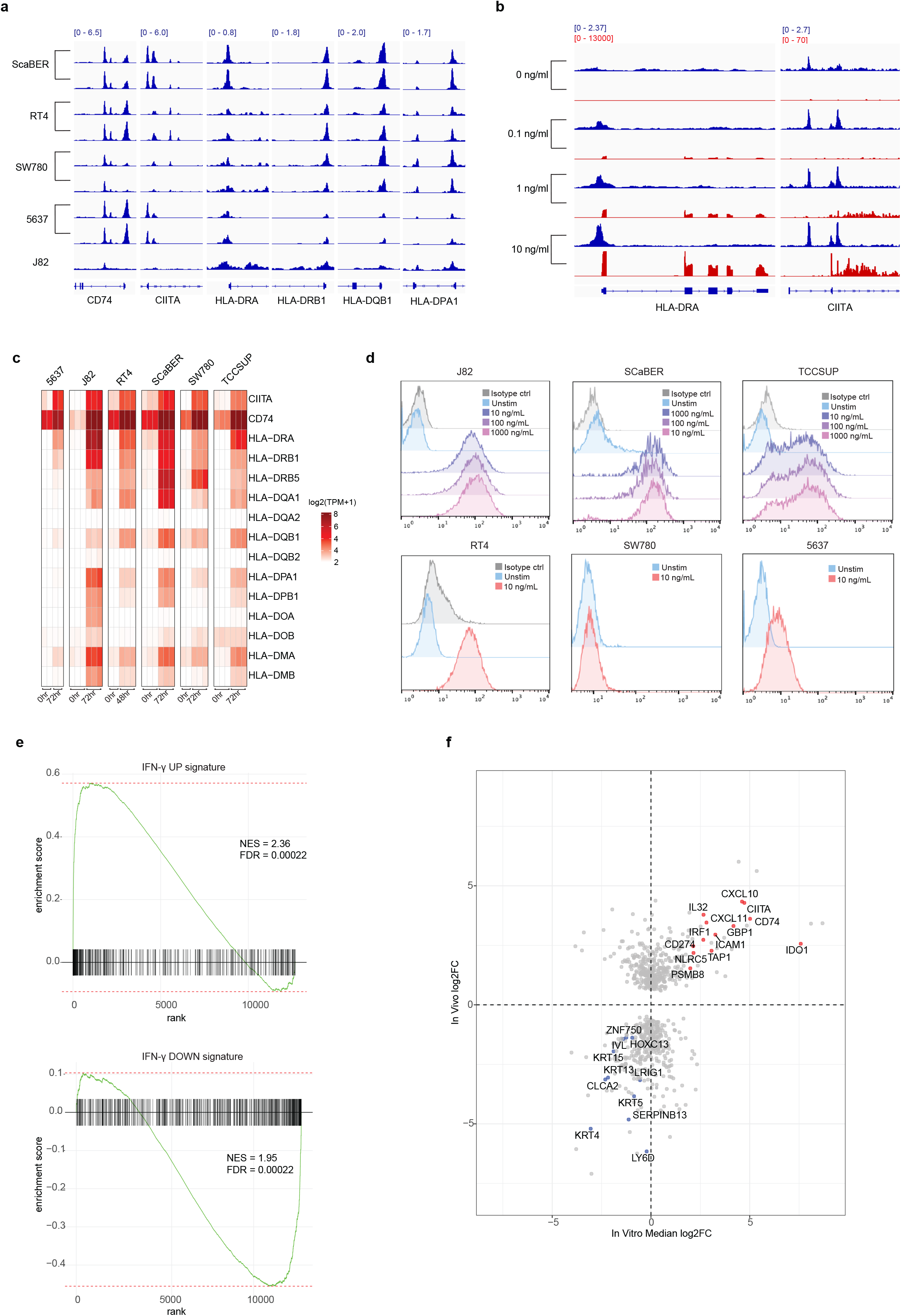
Bladder cancer cells are epigenetically poised for MHC-II expression and respond to IFN-γ with coordinated transcriptional activation and surface protein display. **(a)** Representative Integrative Genomics Viewer (IGV) browser tracks showing baseline ATAC-seq chromatin accessibility near the transcription start sites of six MHC-II loci (CIITA, CD74, HLA-DRA, HLA-DRB1, HLA-DQB1, HLA-DPA1) in five bladder cancer cell lines. Within each gene panel, all tracks share a common y-axis scale (indicated in the upper-left corner); RefSeq gene annotations are shown at the bottom. Most cell lines display accessible chromatin at MHC-II promoters at baseline despite negligible transcriptional output, indicating an epigenetically poised state. J82 and SW-780 are exceptions, showing minimal baseline accessibility. **(b)** IGV browser tracks showing ATAC-seq chromatin accessibility (blue) and RNA-seq signal (red) at MHC-II loci in J82 at baseline and following IFN-γ stimulation. IFN-γ induces de novo chromatin opening at the CIITA and HLA-DRA promoters accompanied by transcriptional activation, demonstrating that inflammatory signaling can actively remodel previously inaccessible MHC-II loci. Shared y-axis scales within each data type are indicated. **(c)** Heatmap of RNA-seq expression values (TPM) for MHC-II pathway genes across six bladder cancer cell lines (SCaBER, TCCSUP, RT4, 5637, SW-780, J82) at 0 hr and 72 hr (48 hr for RT4) following IFN-γ stimulation (10 ng/mL). Five of six lines show robust, coordinated MHC-II induction; SW-780 shows an attenuated transcriptional response. **(d)** Representative flow cytometry histograms showing surface MHC-II (HLA-DR) protein expression in six bladder cancer cell lines at baseline (gray/unfilled) and after 72 h IFN-γ stimulation at the indicated dose (colored/filled). Five of six lines display clear surface MHC-II upregulation. SW-780 is the sole non-responder at the protein level, consistent with the locus-specific chromatin barrier shown in Supplementary Figure 2f. **(e)** Gene set enrichment analysis (GSEA) testing the consensus in vitro IFN-γ response signature in patient tumor cells. The in vitro IFN-γ UP signature is enriched in MHC-II-high malignant epithelial cells, and the in vitro IFN-γ DOWN signature is enriched in MHC-II-low cells (internal snRNA-seq cohort). Gene sets are ranked by normalized enrichment score. **(f)** Genome-wide concordance between the in vitro and in vivo IFN-γ responses. Each point is a gene; axes show log₂ fold change in IFN-γ-stimulated versus unstimulated cell lines (median across five responsive lines) and in MHC-II-high versus MHC-II-low malignant cells in vivo (Spearman ρ = 0.27, p < 0.001). MHC-I and MHC-II antigen-presentation genes and interferon-responsive effector genes (CXCL10, CXCL11, IL32, ICAM1) are among the most concordant upregulated features; basal/squamous markers (KRT5, KRT13, ZNF750, IVL) are among the most concordant downregulated features.

To determine whether this pattern extends to primary tumors, we examined tumor-cell tracks from scATAC-seq of two NMIBC samples(21) and nine TCGA MIBC samples(29). Most tumors across disease stages showed accessible chromatin at MHC-II promoters, mirroring the cell lines (Supplementary Figure 2a). Reduced accessibility was confined to a minority, indicating that epigenetic silencing occurs in only a small fraction of bladder cancers in vivo.

Despite widespread accessibility, canonical MHC-II transcription was negligible across cell lines at baseline. RNA-seq showed minimal HLA-DRA, HLA-DRB1, and related gene expression, consistent with an independent panel of 30 bladder cancer cell lines(30) (Supplementary Figure 2b). Some models expressed CD74, but not a coordinated MHC-II complex, indicating transcriptional silence despite chromatin permissiveness. Among the nine TCGA MIBC tumors with matched scATAC and bulk GSVA data, cases with closed MHC-II chromatin were MHC-II-low (Supplementary Figure 2a, Figure 1a), linking accessibility to pathway activity in patients.

#### IFN-γ converts poised chromatin into coordinated MHC-II transcription and surface protein expression

We characterized the dose and time dependence of MHC-II induction in J82 cells. At 72 hours, IFN-γ elicited a dose-dependent transcriptional response that plateaued at approximately 10 ng/mL; at that concentration, activation was evident by 24 hours and sustained through at least 72 hours (Supplementary Figure 2c-e). We therefore analyzed all cell lines after 72 hours with 10 ng/mL IFN-γ.

In the two cell lines lacking baseline MHC-II chromatin accessibility, IFN-γ produced divergent effects. In J82, it increased accessibility at MHC-II regulatory loci in a time- and dose-dependent manner, accompanied by greater transcription, demonstrating remodeling of previously inaccessible promoters (Figure 2b). SW-780, by contrast, failed to open CIITA and HLA-DR promoters and showed only trace transcript induction (Figure 2b; Supplementary Figure 2f). SW-780 nevertheless mounted a broader, albeit weaker, IFN-γ transcriptional response by GSEA, indicating a locus-specific chromatin block rather than global failure of IFN-γ sensing (Supplementary Figure 2g). All remaining lines showed coordinated MHC-II induction with robust IFN-γ and JAK/STAT enrichment; J82, SCaBER, and TCCSUP also showed TNF-α pathway enrichment (Figure 2c; Supplementary Figure 2g).

Flow cytometry confirmed surface MHC-II induction in five of six cell lines after IFN-γ stimulation (Figure 2d). SW-780 was the exception: despite an attenuated but intact broader IFN-γ response, it showed neither de novo accessibility at CIITA and MHC-II loci nor surface MHC-II. These data show that bladder cancer cells are commonly epigenetically poised but transcriptionally quiescent for MHC-II, and that IFN-γ can license functional tumor-cell antigen presentation by converting latent accessibility into coordinated transcription and surface protein expression.

#### MHC-II-positive tumor cells in patients carry the same IFN-γ-induced program

To quantify the concordance between our in vitro and in vivo findings, we derived a consensus IFN-γ response signature by computing the median log₂ fold change across the five responsive cell lines for each gene, combining p-values using Fisher’s method (Methods). We then tested whether this signature was enriched among differentially expressed genes in MHC-II-positive versus MHC-II-negative tumor cells from our snRNA-seq cohort. The in vitro IFN-γ UP signature was strongly enriched in MHC-II-positive tumor cells in vivo (NES = 2.4, FDR < 0.001; Figure 2e), confirming that the transcriptional program induced by IFN-γ in cell lines closely mirrors the molecular state of MHC-II-positive tumors in patients. A complementary genome-wide analysis comparing gene-level fold changes between the two settings revealed significant concordance across thousands of genes (Spearman ρ = 0.27, p < 0.001; Figure 2f), with MHC-I and MHC-II antigen presentation genes among the most concordant upregulated features.

Conversely, the in vitro IFN-γ DOWN signature was strongly enriched in MHC-II-negative tumor cells in vivo (NES = 1.95, FDR < 0.001; Figure 2e). Genes concordantly downregulated in both settings were dominated by basal and squamous differentiation markers, including KRT5, KRT13, ZNF750, and IVL (Figure 2f). This reciprocal pattern links the IFN-γ/MHC-II axis to both tumor-intrinsic antigen presentation and a shift away from the basal/squamous state. Beyond canonical MHC-II components, MHC-II-positive tumor cells expressed interferon-responsive CXCL10, CXCL11, IL32, and ICAM1, which support immune-cell recruitment and adhesion and extend the program beyond antigen presentation (Figure 2f; Supplementary Figure 2h).

#### IFN-γ induces tumor-cell MHC-II in primary human tumors through JAK/STAT signaling

To assess IFN-γ–mediated MHC-II induction directly in primary human tumors, freshly resected bladder tumors from three patients were dissociated and cultured ex vivo with IFN-γ, with or without the JAK1/2 inhibitor ruxolitinib. Analysis was restricted to the CD45-compartment, isolating tumor cells from infiltrating immune populations (Supplementary Figure 2i). The three tumors revealed two regulatory modes. Two were largely HLA-DR–negative at baseline (patients 1 and 4); in both, IFN-γ induced surface HLA-DR on a subset of tumor cells, and this induction was abolished by ruxolitinib, showing that IFN-γ activates the tumor MHC-II program through canonical JAK/STAT signaling, as in our cell lines. The third tumor (patient 3) showed high baseline HLA-DR that did not increase further with IFN-γ and was not reduced by ruxolitinib, consistent with constitutive, IFN-γ–independent expression. This constitutive state was not seen in our cell-line panel, which was uniformly silent at baseline, and identifies a second route to tumor MHC-II positivity in patients. Although limited to three tumors, these data show that tumor-cell MHC-II is IFN-γ– and JAK/STAT–dependent in baseline-low tumors and can be maintained independently of acute IFN-γ in tumors that already express it, raising the question of what supplies IFN-γ within the bladder tumor microenvironment.

### A discrete IFN-γ–producing, heat-shock–enriched CD8 T-cell subset is present in bladder tumors

In melanoma, IFN-γ produced by tumor-infiltrating T cells drives CIITA-dependent MHC-II induction, linking immune activation to enhanced tumor immunogenicity(13,17,18). Similar inflammatory programs correlate with immune infiltration and response to checkpoint blockade in urothelial carcinoma(19). We therefore asked whether a comparable T cell-driven IFN-γ circuit operates in MHC-II-positive bladder cancers. *IFN-γ production is concentrated in a heat-shock-enriched T-cell population*

Both NMIBC and MIBC scRNA-seq cohorts showed greater T-cell and B-cell infiltration and reduced fibroblast content in MHC-II-high tumors (Supplementary Figure 3a). Because tumor cells comprised approximately 86% of nuclei, however, these datasets contained too few immune cells to resolve the source of IFN-γ. We therefore assembled a complementary TME-enriched cohort from fresh biopsies and cystectomy specimens, yielding greater than 90% viable TME cells and 34,703 sequenced cells across 16 samples (Supplementary Figure 3b,c; Methods). In this cohort, IFN-γ expression localized predominantly to T cells (Supplementary Figure 3d).

Sub-clustering of 23,181 T cells (Figure 3a) and annotation with canonical markers (Figure 3b; Methods) identified a discrete subset characterized by high *IFNG* expression (Figure 3c) together with co-expression of stress-associated chaperone genes (*HSPA1A*, *HSPA1B*, *HSPA6*; Figure 3d). This population was the predominant source of IFN-γ within the T-cell compartment, with the only other appreciable *IFNG* signal coming from exhausted T cells (Tex; Figure 3b,c). Notably, it expressed relatively low levels of both *CD8A* and *CD4* (Figure 3b), raising questions about its precise lineage identity. We refer to this population as Thsp (T heat-shock protein–enriched) cells.

**Figure 3.**
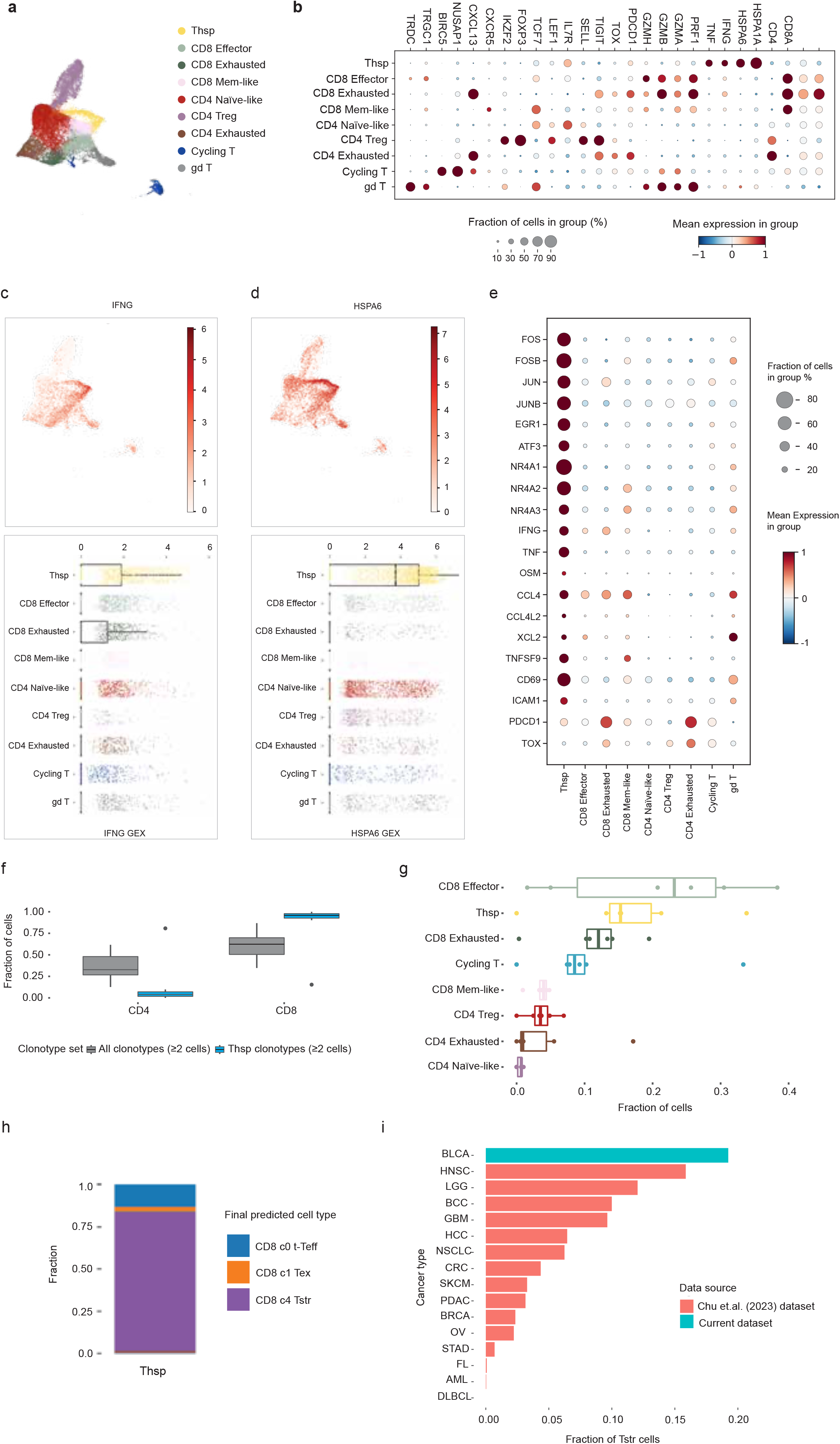
A discrete IFN-γ–producing, heat-shock–enriched T-cell subset (Thsp) is present in bladder tumors. **(a)** UMAP visualization of 23,181 T cells from the TME-enriched scRNA-seq cohort (n = 16 samples, 34,703 total cells), colored by annotated T-cell subtype. Sub-clustering was performed with Seurat and subtypes were assigned based on canonical marker gene expression (Methods). The Thsp population is identified as a discrete cluster. **(b)** Dot plot of canonical marker gene expression (columns) across annotated T-cell subtypes (rows) in the TME-enriched cohort, used to assign subtype identities. Markers span T-cell lineage (CD3D, CD3E, CD4, CD8A), γδ T cells (TRDC, TRGC1), cycling cells (BIRC5, NUSAP1), regulatory T cells (FOXP3, IKZF2), naive and memory states (TCF7, LEF1, IL7R, SELL), cytotoxic effectors (GZMA, GZMB, GZMH, PRF1), and exhaustion (PDCD1, TOX, TIGIT, CXCL13, CXCR5). Dot color represents mean normalized expression (red = higher); dot size indicates the percentage of cells within each subtype expressing the gene. The Thsp population is distinguished by high IFNG and heat-shock chaperone (HSPA1A, HSPA6) expression, lineage-discordant low expression of both CD8A and CD4, and low expression of the exhaustion markers PDCD1 and TOX. **(c)** UMAP of T cells colored by *IFNG* expression (top), with box plot of *IFNG* expression across annotated T-cell subtypes (bottom). *IFNG* expression concentrates within the Thsp cluster. **(d)** UMAP of T cells colored by *HSPA6* expression (top), with box plot of *HSPA6* expression across annotated T-cell subtypes (bottom). *HSPA6* expression concentrates within the same UMAP territory as *IFNG*, identifying Thsp as the predominant intratumoral source of IFN-γ within the T-cell compartment. **(e)** Dot plot of differentially expressed genes (rows) that define the Thsp population, shown across annotated T-cell subtypes (columns) in the TME-enriched cohort. Genes were identified by differential expression of Thsp cells against all other T cells (Methods) and fall into three groups: immediate-early activation genes (FOS, FOSB, JUN, JUNB, EGR1, ATF3) and NR4A family members (NR4A1–3); effector cytokines (IFNG, TNF, OSM) and chemokines (CCL4, CCL4L2, XCL2); and markers of costimulation, retention, and adhesion (TNFSF9, CD69, ICAM1). The exhaustion markers PDCD1 and TOX are shown for contrast. Dot color represents mean normalized expression (red = higher); dot size indicates the percentage of cells within each subtype expressing the gene. The Thsp column shows high expression of the activation and effector program together with low PDCD1 and TOX, defining a non-exhausted effector state. **(f)** TCR clonotype lineage analysis across 14 samples from eight MIBC patients with matched scRNA-seq and scTCR-seq. Points are individual samples. Bars show the fraction of clonotyped cells annotated as CD8 subsets (CD8 Effector, CD8 Exhausted, CD8 Mem-like) or CD4 subsets (CD4 Exhausted, CD4 Naïve-like, CD4 Treg), computed across all clonotypes containing at least two cells (gray) and across the subset of those clonotypes containing at least one Thsp cell (blue). Thsp cells are excluded from both numerators. CD8 subsets are over-represented and CD4 subsets are depleted in Thsp-containing clonotypes (paired Wilcoxon signed-rank test, FDR < 0.025). Per-sample values are shown in Supplementary Figure 3l. **(g)** Fraction of cells in each T-cell subset belonging to a TCR clone (defined as ≥2 cells) that expanded from pre- to post-NAC. Points are the six patients with paired pre- and post-NAC sampling. Subsets with fewer than 50 clonotyped cells in a given patient were excluded. CD8 Effector, Thsp, and CD8 Exhausted subsets show the highest fractions of cells in expanded clones, above all CD4 subsets. **(h)** Proportional bar plots showing the assignment of our T-cell annotations to Chu et al. CD8 atlas annotations (Methods). Approximately 85% of Thsp cells map to the Tstr (stress-response) state defined by Chu et al., and approximately 65% of Tstr cells map to Thsp, indicating substantially overlapping populations. **(i)** Cross-cohort comparison of Tstr abundance across cancer types. CD8 T cells from our bladder cancer cohort (treatment-naive, n = 16) were mapped onto the Chu CD8 atlas using their published reference (Methods) and annotated as CD8 states using the Chu atlas annotation. Bars show the mean fraction of CD8 T cells annotated as Tstr per cancer type across the 15 tumor types in the Chu atlas, with bladder cancer added as a 16th. Bladder cancer ranks first at 19%, compared with 15% in the next-highest tumor type.

Compared with all other T cells, Thsp cells expressed immediate-early activation genes (FOS, FOSB, JUN, JUNB, EGR1, and ATF3) and NR4A-family members, consistent with recent TCR signaling (Figure 3e). They also expressed effector cytokines and chemokines that support lymphocyte recruitment and cDC1 engagement, together with TNFSF9, CD69, and ICAM1, indicating costimulation and retention within the tumor niche. Canonical exhaustion markers PDCD1 and TOX were low. Together, these features define Thsp as an activated, non-exhausted effector population and prompted us to determine its lineage.

#### Thsp cells are clonally expanded CD8 effectors rather than bystander stressed cells

To address the lineage relationships and clonal dynamics of Thsp cells, we analyzed matched scRNA-seq and scTCR-seq data from eight MIBC patients, yielding 14 samples. Six patients were sampled before and after neoadjuvant chemotherapy (NAC) and two contributed a single specimen. The annotated T-cell architecture in these samples recapitulated that of the larger TME-enriched cohort, including the IFN-γ–high, heat-shock–high Thsp population (Supplementary Figure 3e,f,g).

To define lineage, we compared all TCR clonotypes containing at least two cells with the subset that also contained at least one Thsp cell, excluding the Thsp cells themselves from both counts (Methods). CD8 subsets accounted for a median of 58% of clonotyped cells overall and 95% of cells in Thsp-containing clonotypes, whereas the CD4 fraction fell from 40% to 3% (paired Wilcoxon signed-rank FDR < 0.025; Figure 3f). The shift occurred in 13 of 14 samples (Supplementary Figure 3l), placing Thsp within the CD8 lineage despite its low CD8A transcript.

In the six patients with paired sampling, we next calculated for each subset the fraction of cells in clones that expanded after neoadjuvant chemotherapy (Methods). CD8 Effector, Thsp, and CD8 Exhausted cells showed larger expanded-clone fractions than any CD4 subset (Figure 3g). Because intratumoral clonal expansion is a marker of antigen engagement(31,32), these findings, together with the clonotype-sharing analysis, place Thsp within the CD8 compartment responding to tumor antigen rather than a bystander stress state.

Thsp also overlapped a recently described pan-cancer heat-shock-enriched T-cell state, Tstr(33). Projection onto the Chu et al. CD8 atlas mapped approximately 85% of Thsp cells to Tstr, and approximately 65% of cells assigned to Tstr were annotated as Thsp (Figure 3h; Supplementary Figure 3h-k). Thsp and Tstr are therefore largely overlapping populations, with Thsp corresponding to the IFN-γ-producing majority of the stress-response compartment in bladder cancer. Notably, the adverse checkpoint-response association reported for Tstr in other tumor types was absent in the bladder cohort analyzed by Chu et al.

Finally, we mapped all CD8 T cells from our bladder cancer cohort (treatment-naive, n = 16) onto the Chu CD8 atlas using their published reference (Methods) and annotated using their nomenclature. Comparing the mean fraction of CD8 T cells annotated as Tstr per cancer type, bladder cancer showed the highest fraction at 19%, compared with 15% in the next-highest tumor type (Figure 3i).

Together, these data identify Thsp as a discrete, IFN-γ–producing, CD8-lineage T-cell subset that is nested within the broader Tstr population and present across bladder cancer stages. Thsp cells combine recent TCR engagement, inflammatory cytokine and chemokine output, tissue retention, and stress adaptation, without acquiring terminal exhaustion. Together with our tumor-cell findings, these properties position Thsp as a candidate upstream driver of the IFN-γ–licensed tsMHC-II program, raising the question of whether tumors harboring this coordinated program derive greater benefit from immunotherapy.

### tsMHC-II stratifies outcomes across the settings in which bladder cancer is treated with immunotherapy

#### tsMHC-II stratifies clinical outcomes within the luminal NMIBC subtype

We first asked whether tsMHC-II status was prognostic in NMIBC. In the UROMOL cohort, 475 patients had tumor RNA-seq data available; 434 had complete outcome data and were included in survival analyses. In the full analytic cohort, luminal tumors had worse progression-free survival than basal tumors (log-rank p = 0.006; Figure 4a, left). Joint stratification by subtype and MHC-II status localized this difference more specifically: basal/MHC-II-high, basal/MHC-II-low, and luminal/MHC-II-high tumors had similar favorable outcomes, whereas luminal/MHC-II-low tumors had substantially worse progression-free survival (overall log-rank p = 2.3 × 10-4; Figure 4a, right).

**Figure 4.**
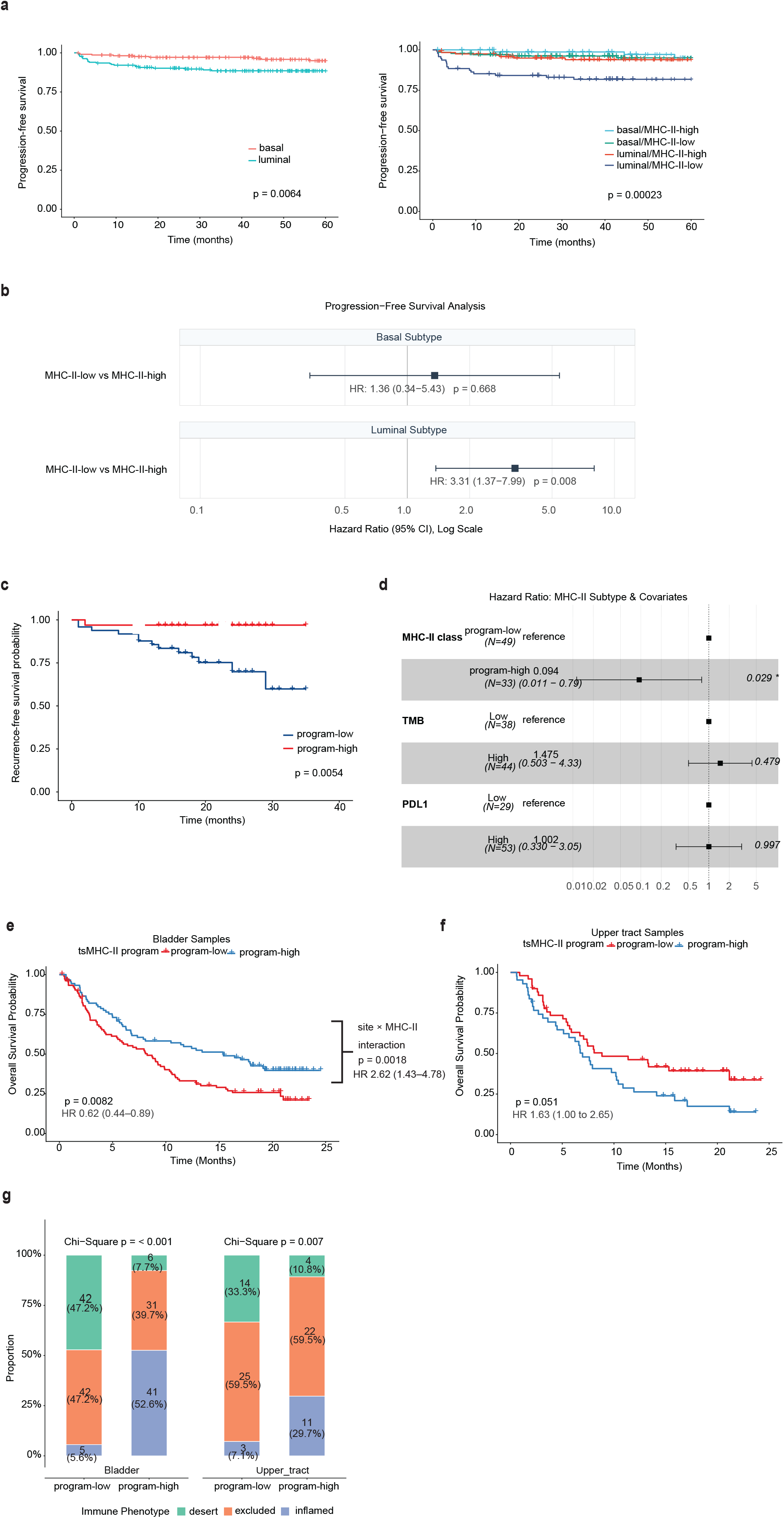
tsMHC-II stratifies outcome within luminal NMIBC and after checkpoint blockade in muscle-invasive and bladder-primary metastatic disease. **(a)** Kaplan-Meier curves for progression-free survival in the UROMOL NMIBC cohort (n = 434 with complete outcome data, predominantly BCG-treated). Left: unadjusted comparison of luminal (n = 216) and basal (n = 218) tumors; log-rank p = 0.006. Right: the same cohort stratified jointly by molecular subtype and MHC-II status, yielding four groups (basal/MHC-II-high, basal/MHC-II-low, luminal/MHC-II-high, luminal/MHC-II-low); overall log-rank p = 2.3 × 10-4. Basal/MHC-II-high, basal/MHC-II-low, and luminal/MHC-II-high tumors show similar and favorable outcomes throughout follow-up. Luminal/MHC-II-low tumors are the sole group with substantially worse progression-free survival, indicating that the subtype-based prognostic difference visible in the left panel is attributable to the enrichment of MHC-II-low tumors within the luminal subtype. **(b)** Subtype-stratified forest plot showing the association between MHC-II-low status and progression-free survival in the UROMOL cohort, estimated separately within luminal (n = 216, 24 progression events) and basal (n = 218, 9 progression events) tumors. Point estimates are hazard ratios from Cox proportional hazards models fitted within each subtype, with MHC-II-high as the reference category; horizontal bars show 95% confidence intervals on a log scale. MHC-II-low status is associated with a 3.3-fold higher risk of progression in luminal tumors (HR 3.31, 95% CI 1.37 to 7.99, p = 0.008) but not in basal tumors (HR 1.36, 95% CI 0.34 to 5.43, p = 0.668). **(c)** Kaplan–Meier survival curves comparing tsMHC-II program–high (n = 33) and tsMHC-II program–low (n = 49) groups in the PURE-01 MIBC cohort, which evaluated neoadjuvant pembrolizumab prior to radical cystectomy. The y-axis shows recurrence-free survival probability; the x-axis shows time. Log-rank test p-value (p = 0.0054) is indicated. MHC-II–high tumors demonstrate markedly improved survival following PD-1 blockade. **(d)** Forest plot showing the results of a multivariate analysis of the PURE-01 MIBC cohort. Group sizes and hazard ratios are indicated in the plot. Only tsMHC-II program-high status shows independent contribution to predicting favorable outcome. Neither TMB nor PD-L1 reached significance in this model. **(e)** Kaplan-Meier curves for overall survival in bladder-primary tumors from IMvigor210 (n = 195), patients treated with atezolizumab, stratified by tsMHC-II program status (program-low, n = 106; program-high, n = 89). Log-rank p = 0.0082. Hazard ratio 0.62 (95% CI 0.44 to 0.89, p = 0.0089) for program-high versus program-low, with program-low as the reference category. The site-by-program interaction test comparing this panel with panel f is shown between the two panels; the interaction term is a ratio of hazard ratios and is not directionally comparable to the within-site estimates. **(f)** Kaplan-Meier curves for overall survival in upper tract primary tumors from the same cohort (n = 93; program-low, n = 50; program-high, n = 43), on identical axes to panel e. Log-rank p = 0.051. Hazard ratio 1.63 (95% CI 1.00 to 2.65) for program-high versus program-low, with program-low as the reference category. Program status was not associated with improved survival at this site. Fewer than 10 patients remain at risk in each group beyond 20 months, so the tails of both curves are unstable. **(g)** Immune phenotype composition by tsMHC-II program status, shown separately for bladder-primary and upper tract primary tumors. Bars show the proportion of tumors classified as desert, excluded, or inflamed, with counts and percentages overlaid. Chi-square p < 0.001 for bladder and p = 0.007 for upper tract. Immune phenotype calls were available for 167 of 195 bladder and 79 of 93 upper tract patients. The inflamed fraction increases and the desert fraction decreases with program-high status at both sites. The excluded fraction differs, it declines with program status in bladder tumors (47.2% to 39.7%) and is unchanged in upper tract tumors (59.5% in both groups).

In subtype-stratified Cox models, MHC-II-low status was associated with greater progression risk in luminal tumors (n = 216, 24 progression events; HR 3.31, 95% CI 1.37 to 7.99, p = 0.008; Figure 4b). The basal estimate was imprecise and not significant (n = 218, 9 progression events; HR 1.36, 95% CI 0.34 to 5.43, p = 0.668; Figure 4b), and the formal subtype-by-MHC-II interaction did not reach significance (p = 0.263). MHC-II status alone produced only marginal separation in the full cohort (log-rank p = 0.039; Supplementary Figure 4a). A main-effects model including subtype and MHC-II status is shown in Supplementary Figure 4b. In a clinically adjusted analysis of the full cohort, MHC-II status did not retain an independent association, whereas tumor grade was the only significant adjusted covariate. These results support a context-dependent prognostic association within luminal NMIBC rather than an independent full-cohort marker.

Whole-genome doubling (WGD) is a recently identified adverse prognostic factor in NMIBC and has been linked to immune evasion(34–36), raising the possibility that tsMHC-II status is a readout of genomic instability rather than an independent axis of immune engagement. We therefore tested whether the two features coincide within UROMOL luminal tumors. Within UROMOL luminal tumors, MHC-II-high cases were not significantly enriched by WGD status (WGD-positive: 50.0% (8/16); WGD-negative: 63.5% (54/85); p = 0.40, Fisher’s exact test; Supplementary Figure 4c). Combined stratification by WGD and MHC-II status yielded four distinct survival trajectories, although the WGD-defined groups were small (n = 8 each for WGD/MHC-II-high and WGD/MHC-II-low) and should be interpreted cautiously. Thus, tsMHC-II provides prognostic stratification within luminal NMIBC that is not explained by enrichment for WGD. Because this cohort does not provide a treatment-comparator analysis, these findings do not establish tsMHC-II as a predictor of BCG benefit.

#### The tsMHC-II program is associated with pathological response and recurrence-free survival after neoadjuvant pembrolizumab

We next asked whether the tumor IFN-γ-responsive state marked by tsMHC-II was associated with outcome after neoadjuvant checkpoint blockade. tsMHC-II-positive tumors expressed a coordinated inflammatory program extending beyond antigen presentation to chemokines, ICAM1, IL32, and additional interferon-stimulated genes. We therefore derived a broader tumor-cell program intended to be more robust than a pure MHC-II gene set in bulk clinical specimens with heterogeneous immune content.

The signature was constructed without reference to clinical outcome by intersecting genes induced by IFN-γ in bladder cancer cell lines with genes enriched in tsMHC-II-positive malignant epithelial cells in the internal NMIBC and external MIBC single-cell cohorts (Methods). The resulting 11 genes span antigen presentation (CD74 and CIITA), T-cell recruitment (CXCL10 and CXCL11), adhesion (ICAM1), and additional interferon-responsive genes (GBP5, IL32, SOD2, TNFRSF1B, HAPLN3, and SLC12A7; Supplementary Table 6). We refer to this set as the tsMHC-II program signature. GSVA enrichment was analyzed continuously where specified and classified as program-high or program-low using outcome-independent, cohort-specific Gaussian mixture cutpoints (Methods).

We applied the signature to PURE-01 (NCT02736266, n = 82), a single-arm phase II trial of neoadjuvant pembrolizumab before radical cystectomy(37). The score showed a bimodal distribution, and the cohort-specific mixture cutpoint classified 33/82 (40%) tumors as program-high (Supplementary Figure 4d). Pathological complete response (ypT0) occurred in 52% (17/33) of program-high versus 24% (12/49) of program-low tumors (OR 3.27, 95% CI 1.15 to 9.11, p = 0.018). Program-high status was also associated with longer recurrence-free survival (log-rank p = 0.0054; Figure 4c). In a multivariable Cox model containing tumor mutational burden and PD-L1 expression (n = 82, 14 events), program-high status retained an independent association with outcome (HR 0.094, 95% CI 0.011 to 0.79, p = 0.029; Figure 4d), whereas TMB and PD-L1 were not significant.

Because MHC-II is also expressed by stromal and immune cells, we tested whether tumor purity accounted for the bulk program association. The score was inversely correlated with tumor purity (Pearson r = -0.62, p < 0.001; Supplementary Figure 4e). After residualizing the score for purity, program status remained associated with recurrence-free survival (HR 0.18, 95% CI 0.05 to 0.65, p = 0.009; Supplementary Figure 4f,g), including in a model containing TMB and PD-L1 (HR 0.14, 95% CI 0.03 to 0.73, p = 0.019; Supplementary Figure 4h). Thus, tumor purity explains part of the score variance but not the observed survival association. Because PURE-01 is single-arm, these findings establish association with outcome after pembrolizumab, not treatment-predictive value.

#### The tsMHC-II program is associated with survival in bladder-primary metastatic urothelial carcinoma

We next evaluated the tsMHC-II program in IMvigor210, a single-arm phase II trial of atezolizumab in platinum-treated locally advanced or metastatic urothelial carcinoma. Of 348 patients with pretreatment expression and overall survival data, 8 lacked a reported tissue of origin and 52 were profiled from a non-primary specimen. We restricted analysis to 288 primary-site specimens because bulk MHC-II signal from lymph nodes and other non-primary tissues can reflect resident antigen-presenting cells (Supplementary Figure 5a; Methods). The analytic cohort comprised 195 bladder-primary and 93 upper tract tumors. A cohort-specific Gaussian mixture cutpoint classified 89/195 (45.6%) bladder and 43/93 (46.2%) upper tract tumors as program-high (Supplementary Figure 5b; Supplementary Table 7). Bladder-primary and upper tract tumors were comparable in sex, performance status, tumor mutational burden, PD-L1 immune-cell level, and both Lund and TCGA subtype composition, and differed materially only in prior platinum exposure (Supplementary Table 8). Separate analysis by primary site was post hoc; the pooled analysis is shown in Supplementary Figure 5c.

In bladder-primary tumors, program-high status was associated with longer overall survival (log-rank p = 0.0082; HR 0.62, 95% CI 0.44 to 0.89, p = 0.0087; Figure 4e). The continuous score gave a concordant result (HR 0.82 per standard deviation, p = 0.026). Among the 177 patients with complete clinical covariate data, the unadjusted HR was 0.66 (95% CI 0.46 to 0.95); adjustment for ECOG performance status, liver metastasis, prior platinum therapy, and sex attenuated the estimate to 0.76 (95% CI 0.52 to 1.10, p = 0.143; Supplementary Figure 5d), indicating that clinical prognostic factors accounted for part of the association despite being closely balanced between program groups (Supplementary Table 7). Among 157 patients with evaluable TMB, the association persisted after TMB adjustment (HR 0.59, 95% CI 0.39 to 0.90, p = 0.014). Adding PD-L1 immune-cell staining produced a similar effect estimate with wider uncertainty (HR 0.64, 95% CI 0.40 to 1.02, p = 0.061; Supplementary Figure 5d).

In upper tract tumors, program-high status was not associated with improved overall survival (log-rank p = 0.051; HR 1.63, 95% CI 1.00 to 2.65; Figure 4f). The unadjusted site-by-program interaction suggested different associations by primary site (interaction HR 2.62, 95% CI 1.43 to 4.78, p = 0.0018; Supplementary Figure 5e). The interaction remained significant in the complete-case subset (p = 0.021) but attenuated after adjustment for clinical prognostic factors (p = 0.059). No association was detectable when both primary sites were pooled (HR 0.85, 95% CI 0.64 to 1.14, p = 0.277; Supplementary Figure 5c). These findings suggest an anatomic source of heterogeneity that requires confirmation rather than defining a validated boundary for biomarker use.

The program tracked with immune context at both sites. The inflamed fraction increased from 5.6% in program-low to 52.6% in program-high bladder tumors and from 7.1% to 29.7% in upper tract tumors (p < 0.001 and p = 0.007, respectively; Figure 4g). The excluded fraction declined with program status in bladder tumors but remained 59.5% in both upper tract groups, consistent with the T-cell-depleted contexture described for upper tract urothelial carcinoma(38). Objective response was not significantly enriched in program-high bladder tumors (30.4% versus 20.2%, p = 0.154) and was less frequent in program-high upper tract tumors (5.7% versus 26.7%, p = 0.018; Supplementary Figure 5f), although the latter comparison included only 2 responders among 35 program-high patients. The discordance between an inflamed phenotype and response in upper tract disease suggests that the bulk program is not simply an immune-infiltrate readout.

Together, these analyses associate tumor-cell MHC-II with clinically relevant outcomes across bladder cancer stages. In NMIBC, MHC-II status stratified progression risk within luminal tumors. In two independent single-arm checkpoint-treated cohorts, the tumor-cell-derived 11-gene program was associated with pathological response or survival, with the PURE-01 association persisting in a model containing TMB and PD-L1 and the bladder-primary IMvigor210 association persisting after TMB adjustment. These findings support prospective evaluation of the program as a bladder-primary biomarker.

## DISCUSSION

Tumor-cell MHC-II is associated with clinical outcomes at three points in the bladder cancer treatment course, although the evidence differs by setting. Within luminal NMIBC, where BCG is standard of care, MHC-II-low tumors carry the excess progression risk historically attributed to luminal classification as a whole; this is a prognostic observation rather than evidence of BCG-specific benefit. In muscle-invasive disease treated with neoadjuvant pembrolizumab, an 11-gene tumor-cell program is associated with pathological complete response and recurrence-free survival and retains an association in a model containing TMB and PD-L1. In bladder-primary metastatic disease treated with atezolizumab, the same signature is associated with overall survival, persisting after adjustment for TMB but attenuating after adjustment for clinical prognostic factors. This consistency across distinct settings supports prospective validation.

What makes this measurable is that the signal is tumor-cell-derived. Malignant epithelial cells account for the difference in MHC-II pathway activity between high and low tumors, even though professional antigen-presenting cells retain higher per-cell expression, and the proportion of MHC-II-positive cells of tumor origin rises from 26% to 68% between the two groups. This matters practically: a bulk score that tracked infiltrate composition would largely restate what PD-L1 already measures. Approximately one third of bladder tumors carry the phenotype by both tumor-specific measures, single-cell expression and HLA-DR protein staining, a prevalence that corresponds to roughly 26,700 of the approximately 80,000 bladder cancers diagnosed annually in the United States.

The program is inducible rather than fixed, and this determines how a negative result should be read. MHC-II regulatory loci are accessible but transcriptionally silent in most bladder cancer cell lines and in most primary tumors, and IFN-γ converts this poised configuration into coordinated transcription and surface protein through JAK/STAT signaling, as the ex vivo ruxolitinib experiments in primary tumors confirm. Accessibility is therefore necessary but not sufficient, and because it is already present in most tumors it rarely discriminates high from low disease in vivo, where IFN-γ availability appears to be the limiting variable. This competence appears to be a retained property of the urothelial lineage rather than a cancer-acquired trait: in differentiated normal human urothelium from six donors, IFN-γ uniformly induced the MHC-II machinery along with the same CXCR3-ligand chemokines that define our tumor program(39). The clinical consequence is that most MHC-II-low tumors are not incapable of antigen presentation, they are uninstructed. The output of the program extends beyond antigen presentation. MHC-II-positive tumor cells co-express CXCL10, CXCL11, ICAM1, and IL32, which recruit CXCR3⁺ effectors and stabilize immunological synapses. This is why we defined the biomarker on the broader 11-gene program rather than on MHC-II genes alone, and why the phenotype should be understood as tumor-cell participation in an inflamed niche rather than as a single-pathway readout.

We identify Thsp cells, a heat-shock-enriched CD8-lineage subset, as a candidate source of the licensing IFN-γ. These cells combine IFN-γ production with features of recent TCR engagement and tissue retention while lacking canonical exhaustion markers, and clonotype sharing with conventional CD8 effectors together with clonal expansion after neoadjuvant chemotherapy supports antigen-driven activation rather than bystander stress. Approximately 85% of Thsp cells map to the pan-cancer Tstr stress-response state and approximately 65% of Tstr cells map to Thsp, so the two populations largely coincide in bladder cancer, with Thsp marking the IFN-γ-producing fraction. This matters because stress-associated T-cell signatures have been linked to immunotherapy resistance in other tumor types, yet that association was specifically absent in the urothelial cohort tested by Chu and colleagues. Whether the discordance reflects a bladder-specific property of the stress state, or the IFN-γ output that distinguishes Thsp, will require larger single-cell-profiled clinical cohorts.

In NMIBC the reading is prognostic and subtype-dependent. MHC-II status did not retain an independent association in the clinically adjusted full-cohort analysis, in which tumor grade was the only significant adjusted covariate. The more informative observation is that tsMHC-II stratifies within the luminal subtype: luminal MHC-II-high tumors show survival comparable to basal disease, while luminal MHC-II-low tumors carry the excess risk. The nonsignificant formal interaction and imprecise basal estimate limit a claim of strict subtype specificity. Nevertheless, tsMHC-II provides a mechanistic explanation for prognostic heterogeneity within luminal bladder cancer rather than functioning as an orthogonal full-cohort marker. Its lack of enrichment by whole-genome doubling supports a distinct axis of immune engagement, with potential implications for heterogeneity in BCG response.

Two observations qualify interpretation of the biomarker. First, its association with outcome may differ by primary site. The survival association was observed in bladder-primary tumors but not upper-tract tumors, and the site-by-program interaction, although significant before clinical adjustment, attenuated afterward. The immune phenotype data offer one possible explanation: although the program tracked with a shift from immune-desert toward inflamed contexture at both sites, the excluded fraction remained unchanged in upper-tract tumors, consistent with immune recognition that does not translate into corresponding infiltration. Bladder and upper-tract urothelial carcinomas share urothelial histology but differ in developmental and etiologic context, the relative frequency of molecular drivers, and immune contexture(40). Exploratory subgroup analyses from CheckMate 274(41), and AMBASSADOR(42) have also suggested that systemic treatment effects may vary by primary site. Because these upper-tract subgroups were small and not powered for site-specific efficacy, the collective evidence (including our post hoc IMvigor210 analysis) supports primary site as a hypothesis-generating stratification factor rather than a validated predictive biomarker.

The second is the relationship to molecular subtype, which is not constant across stages. In NMIBC the phenotype is enriched within luminal tumors, whereas in metastatic disease program-high tumors are enriched for basal and depleted for UroA classification (Supplementary Table 7), consistent with the basal localization reported by Xiong and colleagues in muscle-invasive disease(20). These observations may reflect stage-dependent differences in inflammatory input, although differences among subtype classifiers cannot be excluded. Xiong and colleagues also reported an inverse survival association in the untreated, mixed-treatment TCGA cohort while their checkpoint-blockade findings align with ours. One contributor may be methodological: they used a survival-optimized cutpoint, whereas our outcome-independent Gaussian mixture cutpoints were estimated separately within each cohort.

Two assay formats are potentially suitable for clinical use. The 11-gene program score can be computed from bulk RNA-seq, and HLA-DR immunohistochemistry with a validated antibody clone and blinded histoscore reading is directly deployable in a diagnostic laboratory, as demonstrated here across 122 specimens. The clinical context for such a biomarker is changing. Enfortumab vedotin plus pembrolizumab is now first-line standard of care in advanced urothelial carcinoma, and our checkpoint-treated cohorts predate this transition. They therefore establish the association of tsMHC-II with outcome after checkpoint blockade rather than its value in contemporary combination therapy. Because tsMHC-II marks an IFN-γ-responsive, immune-engaged state, it may report the immunotherapy-sensitive component of such regimens, and its utility may ultimately emerge in combination with biomarkers of ADC sensitivity rather than as a standalone predictor. Immunotherapy monotherapy also remains standard in the adjuvant setting and in patients unable to receive the combination.

Several limitations should be acknowledged. The checkpoint-blockade cohorts are both single-arm, so these analyses cannot separate prognostic from predictive effects, and demonstrating predictive value will require a randomized or comparator-controlled cohort. The metastatic analysis was restricted to primary-site specimens on measurement-validity grounds and the separate analysis by primary site was post hoc, so the site-specific findings require confirmation. The evidence that Thsp cells are the physiological IFN-γ source in vivo remains correlative; the ruxolitinib experiments establish that IFN-γ drives tumor-cell MHC-II through JAK/STAT signaling, but not that Thsp-derived IFN-γ is necessary or sufficient. Several analyses rest on limited patient numbers, including the single-cell TCR cohort and the ex vivo stimulation experiments.

Finally, prospective validation will be required to establish biomarker utility, to define optimal assay implementation, choose between RNA and protein formats, and define performance in the combination era.

In summary, tumor-cell MHC-II identifies an inducible, IFN-γ-dependent state present in approximately one third of bladder cancers and associated with clinically relevant outcomes from NMIBC through metastatic disease. In checkpoint-treated cohorts, the 11-gene program retained associations in models containing established tissue biomarkers, while its treatment-predictive value remains to be tested in comparator-controlled studies. The state is measurable by two assays compatible with clinical laboratories, although a locked clinical threshold remains to be established. In these tumors, malignant cells are not passive targets but active participants in the immune response. Licensing, unlike mutation, can be supplied, so the value of tsMHC-II lies not only in identifying tumors that already present antigen but also in defining an inducible state that may be amenable to therapeutic conversion.

## METHODS

### Patient cohorts and datasets

#### Internal NMIBC cohort (Bellmunt cohort)

FFPE specimens from non-muscle-invasive bladder cancer patients were obtained from the Hospital del Mar–Parc de Salut Mar Biobank, Barcelona, Spain. Written informed consent was obtained from all patients. Bulk RNA-seq was obtained from GSE136401. FiTAc-seq H3K27ac profiling on 16 specimens (six with a micropapillary component), snRNA-seq on nine specimens, and scATAC-seq on two specimens were obtained from GSE281743. Clinical annotations including treatment history (predominantly BCG) and progression-free survival outcomes were obtained from the biobank registry.

#### Internal TME-enriched scRNA/scTCR-seq cohort

Fresh tumor biopsies were obtained at transurethral resection (TURB) from 16 treatment-naive patients with primary bladder cancer at Germans Trias i Pujol Institute for Health Science Research (IGTP). Two additional MIBC patients had paired pre- and post-neoadjuvant chemotherapy (NAC) sampling and were profiled by combined scRNA-seq and scTCR-seq. The study was approved by the Research Ethics Committee (Reference PI-19-111) and the Institutional Review Board (IRB-00005712). Written informed consent was obtained from all patients.

#### Tumor microarray cohort

IHC was performed on a previously described TMA(25) comprising 162 bladder tumor specimens (102 HGT1, 41 low-grade tumors, and 19 MIBC), of which 122 were evaluable for MHC-II scoring.

#### Public bulk transcriptomic cohorts

The UROMOL cohort (n = 535 NMIBC patients, predominantly BCG-treated) as described by Lindskrog et al. (23) was obtained from EGAD00001006656. The TCGA-BLCA cohort (n = 410 MIBC patients) was downloaded from the Genomic Data Commons. The PURE-01 cohort (n = 82; phase II trial of neoadjuvant pembrolizumab prior to radical cystectomy; NCT02736266) was obtained from EGAD00001008003 as described by Basile et al.(37). The Bellmunt internal NMIBC bulk RNA-seq cohort (n = 62) is described above and in Bowden et al.,(43).

#### Public single-cell and chromatin datasets

The external MIBC scRNA-seq cohort (n = 25 patients, 10x Genomics 3′ chemistry; 67,988 cells after quality filtering) was obtained from GSE169379(24). Nine MIBC scATAC-seq samples were obtained from https://gdc.cancer.gov/about-data/publications/TCGA-ATAC-Seq-2024 (29). The bladder cancer cell line RNA-seq panel (n = 30 lines) was obtained from GEO accession GSE97768(30). The Chu et al. pan-cancer CD8 T-cell reference was obtained from https://singlecell.mdanderson.org/TCM/download/CD8.

#### Clinical endpoint definitions

In the UROMOL NMIBC cohort, the primary endpoint analyzed was progression-free survival (PFS), defined as the interval from diagnosis to progression to muscle-invasive disease or last follow-up. In the PURE-01 MIBC cohort, the primary endpoint analyzed was recurrence-free survival (RFS), defined as the interval from radical cystectomy to recurrence or last follow-up; pathological complete response was defined as the absence of residual invasive carcinoma (ypT0) versus any residual disease at cystectomy.

#### Ethics approval and consent

All procedures involving human participants or human tissue were conducted in accordance with the Declaration of Helsinki and were approved by the relevant institutional research ethics committees. The internal TME-enriched cohort was approved by the Research Ethics Committee (PI-19-111) and Institutional Review Board (IRB-00005712), and all participants provided written informed consent. Analyses of deidentified publicly available datasets were conducted under the approvals and consent provisions reported in the original studies.

### Tissue methods

#### FiTAc-seq H3K27ac profiling

For H3K27ac profiling we selected 16 NMIBC specimens, six of them with a micropapillary component. To increase enrichment for cancer cells, FFPE sections were macrodissected when needed to obtain >80% tumor cells. The FiTAc-seq method was applied as previously described(44). Briefly, 10 sections of 10 μm thickness were washed three times with xylenes to remove paraffin, then rehydrated through an ethanol/water series. Tissue was resuspended in lysis buffer as previously described and sonicated for 5 minutes using a Covaris E220 instrument (setting: 140 peak incident power, 5% duty factor, 200 cycles per burst) in 1 ml adaptive focused acoustics (AFA) fiber millitubes. Soluble chromatin (5 μg) was immunoprecipitated with 10 μg H3K27ac antibody (Diagenode catalog number C15410196). ChIP-seq libraries were constructed using ThruPLEX-FD kits (Rubicon Genomics) following the manufacturer’s protocols. 75-bp single-end reads were sequenced on a NextSeq instrument (Illumina).

#### Nuclei isolation and snRNA-seq from FFPE tissue

Nine NMIBC clinical samples were selected as described above. 10 μm sections were prepared and washed three times with xylenes to remove paraffin, then rehydrated through an ethanol/water series. A modified version of the nuclei isolation protocol for frozen specimens(45) was applied with the following modifications. After paraffin removal and rehydration with a graded alcohol series ending in water, centrifugation, and water removal, tissue was resuspended in a buffer containing 0.1% NP40, 0.1% Tween-20, and 0.01% digitonin. The homogenate was transferred to a pre-chilled 1.5 ml microfuge tube and incubated on ice for 10 min. Lysates were filtered through a 40 μm cell strainer, and nuclei were centrifuged for 10 min at 1500 relative centrifugal force in a pre-chilled (4°C) fixed-angle centrifuge. Nuclei were resuspended in 300 μl of buffer containing 0.1% Tween-20 and enumerated using a hemocytometer with Trypan blue staining. Approximately 10,000 nuclei per sample were fixed and processed using the 10x Genomics Chromium Fixed RNA Profiling (FLEX) kit per manufacturer’s protocol. Libraries were multiplexed and sequenced on a NovaSeq XP instrument with 150 bp paired-end reads to a target depth of 10,000 reads per nucleus.

#### Tissue processing for TME and TCR analysis

Tumor tissue samples were obtained from the exophytic part of the tumor during transurethral resection (TURB) from treatment-naïve patients with primary bladder cancer. Immediately after surgery, biopsies were placed in RPMI 1640 medium (Thermo Fisher Scientific, Spain) and transported to the laboratory. Upon arrival, tissue samples were minced using a sterile scalpel in RPMI medium and processed into single-cell suspensions by enzymatic digestion. Tissue fragments were incubated with Collagenase II (Sigma, Spain; 0.5 mg/mL) and DNase I (1 U/mL) in RPMI supplemented with 5% fetal bovine serum (FBS) at 37°C under continuous agitation. Digestion was performed over two to three successive cycles of 30 min each. The resulting cell suspensions were filtered through a 40 μm cell strainer (Becton & Dickinson) to remove debris, and viable cells were cryopreserved in freezing medium containing dimethyl sulfoxide (DMSO).

#### scRNA-seq and scTCR-seq library preparation (TME-enriched cohort)

Single-cell suspensions from dissociated TURB biopsies (16 patients) were processed using 10x Genomics Chromium Fixed RNA Profiling (FLEX) kit as above. For the two NAC-paired patients (v61, v70), single-cell suspensions were processed using the 10x Genomics 5′ v2 chemistry combined with V(D)J enrichment for paired α/β TCR profiling. Libraries were sequenced on NovaSeq with targeted read depth of 20,000/cell.

#### Immunohistochemistry

IHC was performed on a tissue microarray made from 162 bladder tumor samples (102 HGT1, 41 low-grade, 19 MIBC). Tissue sections were deparaffinized in xylenes and hydrated through ethanol/water series. After antigen retrieval, slides were treated with 3% H₂O₂ in PBS for 10 min to quench endogenous peroxidases, washed, and incubated in blocking solution (PBS containing 1% BSA and 1% Tween-20) for 1 h at ambient temperature. MHC-II staining was performed with anti–HLA-DR antibody (DAKO cat#M0775, clone CR3/43) as previously validated for tumor MHC-II detection(18) diluted in blocking solution and incubated for 1 h. Slides were washed in PBS and incubated with the peroxidase-based EnVision Kit (Dako).

#### Staining scoring

MHC-II staining was assessed by Histoscore (H-score) by an expert genitourinary pathologist blinded to clinical outcome and molecular subtype. H-score was calculated as: (3 × percentage of strongly staining cells) + (2 × percentage of moderately staining cells) + (1 × percentage of weakly staining cells), yielding a range of 0–300. Staining intensity was scored as: 0, no staining; 1, weak; 2, moderate; 3, strong. Tumors were classified as MHC-II–positive when H-score > 2 with membranous staining present on tumor cells as established in prior melanoma studies(18).

#### Ex vivo IFN-γ stimulation and HLA-DR analysis in primary bladder tumors

For ex vivo functional studies, freshly dissociated tumor samples were filtered through a 100 μm cell strainer (BD Biosciences) to preserve larger cellular aggregates and maximize cell recovery. Cells were seeded into 24-well culture plates and maintained overnight in complete RPMI medium at 37°C and 5% CO₂. Twenty-four hours after isolation, cultures were stimulated with recombinant human IFN-γ (Biolegend, Palex, Spain) at final concentrations of 50 ng/mL for 72 hours. To inhibit IFN-γ-induced JAK/STAT signaling, cells were pretreated with the selective JAK1/2 inhibitor ruxolitinib (20 μM) kindly provided by Dr. Ester Ballana (IrsiCaixa Foundation), for 1 hour before cytokine stimulation. Following culture, cells were harvested and stained for flow cytometric analysis. Briefly, cells were incubated with Human TruStain FcX™ Fc Receptor Blocking Solution (BioLegend, Palex, Spain) to minimize nonspecific antibody binding.

Cell viability was assessed using Fixable Viability Stain 575V (1:1500 dilution, 30 min at 4°C; BD Biosciences). Surface staining was performed using anti-CD45 BV421, anti-EpCAM BV605, and anti-HLA-DR BV711 antibodies. After staining, samples were washed and acquired on an LSRFortessa flow cytometer (BD Biosciences) at the Flow Cytometry Core Facility of the Germans Trias i Pujol Research Institute. Data were analyzed using FlowJo software (v10.10; BD/Tree Star, Portland, OR, USA). Tumor cells were identified as viable CD45⁻ cells, and HLA-DR expression was quantified within the tumor-cell compartment.

### Cell-line and in vitro methods

#### Cell lines and culture conditions

The human bladder cancer cell lines J82 (HTB-1), TCCSUP (HTB-5), SCaBER (HTB-3), 5637 (HTB-9), RT4 (HTB-2), and SW780 (CRL-2169) were procured from ATCC and maintained under conditions recommended by ATCC. J82, SCaBER, and TCCSUP were cultured in Eagle’s Minimum Essential Medium (EMEM) supplemented with 10% fetal bovine serum (FBS) and 1% penicillin-streptomycin (Pen-Strep). RT4 and 5637 were grown in RPMI-1640 with 10% FBS and 1% Pen-Strep. SW780 cells were cultured in Leibovitz’s L-15 medium with 10% FBS and 1% Pen-Strep. All cell lines were maintained at 37°C in a humidified incubator with 5% CO₂. Cell lines were tested and found negative for mycoplasma contamination.

#### IFN-γ stimulation

Cells were plated in triplicate on 100 mm dishes at a density of 0.5 × 10⁶ cells per dish and permitted to adhere overnight. The following day, cells were washed to remove dead cells and debris. Recombinant human IFN-γ (BioLegend, catalog 570208) was reconstituted per the manufacturer’s instructions and added directly to the culture medium to reach the indicated final concentration (0.1–1000 ng/mL for dose-response experiments; 10 ng/mL for time-course and standard stimulation experiments). Control (unstimulated) cells received an equal volume of vehicle (PBS). Cells were incubated for the indicated time points (0, 24, 48, or 72 h), then harvested by trypsinization, centrifuged at 800 × g for 5 min, resuspended in Bambanker Cell Freezing Medium, and stored at −80°C until further processing.

#### RNA isolation

Total RNA was extracted using TRIzol Reagent (Thermo Fisher Scientific) per manufacturer’s protocol. Cells were thawed at 37°C and centrifuged at 1,500 × g for 2 min. Pellets were lysed by adding 1 mL TRIzol followed by 0.2 mL chloroform and centrifuged at 12,000 × g for 15 min at 4°C. The aqueous phase was transferred to a fresh RNase-free tube and RNA precipitated by adding 0.5 mL isopropanol per 1 mL TRIzol used. Samples were centrifuged at 12,000 × g for 10 min and the RNA pellet washed with 1 mL of 75% ethanol, air-dried on ice, and resuspended in 20 μL of RNase-free water. RNA concentration and purity were assessed on a Nanodrop spectrophotometer (Thermo Fisher Scientific). Samples were stored at −80°C until library preparation.

#### RNA library preparation and sequencing

Library preparation and RNA sequencing were performed at the Molecular Biology Core Facility (MBCF) at Dana-Farber Cancer Institute. RNA integrity was confirmed by Bioanalyzer (RIN ≥ 8). Strand-specific libraries were prepared using ribosomal RNA depletion, RNA fragmentation, synthesis of first- and second-strand cDNA, adapter ligation, PCR amplification, and barcode application. Libraries were sequenced on a NovaSeq X-Plus instrument to obtain approximately 50 million 150-bp paired-end reads per sample.

#### ATAC-seq library preparation

The ATAC-seq protocol was adapted from the Omni-ATAC protocol(46). Viably frozen cells were thawed at 37°C, pelleted by centrifugation, and counts and viability were assessed using a Countess cell counter with NucBlue stain. 200,000 cells were resuspended in 1 mL of cold ATAC-seq resuspension buffer (RSB; 10 mM Tris-HCl pH 7.4, 10 mM NaCl, 3 mM MgCl₂ in water). Cells were centrifuged at 800 × g for 5 min in a pre-chilled (4°C) fixed-angle centrifuge and the supernatant carefully aspirated. Pellets were resuspended in 50 μL of ATAC-seq RSB containing 0.1% NP40, 0.1% Tween-20, and 0.01% digitonin by gentle pipetting and incubated on ice for 3 min. Lysis was stopped by addition of 1 mL of ATAC-seq RSB containing 0.1% Tween-20 (without NP40 or digitonin), and tubes were inverted to mix. Nuclei were centrifuged for 10 min at 800 × g (4°C). Supernatant was removed and nuclei were resuspended in 50 μL of transposition mix (25 μL 2× TD buffer, 2.5 μL transposase Tn5 [Illumina 20034197], 16.5 μL PBS, 0.5 μL 1% digitonin, 0.5 μL 10% Tween-20, 5 μL water) by gentle pipetting. Transposition was performed at 37°C for 30 min in a thermomixer at 1,000 rpm. Reactions were cleaned up with the Qiagen MinElute kit. Libraries were amplified with qPCR optimization as described [Buenrostro et al., *Curr Protoc Mol Biol*, 2015], purified with Qiagen QIAquick PCR purification columns, and quantified using Qubit. Libraries were sequenced on an Illumina Novoseq X plus with paired-end 150 base pair reads with a targeted depth of 30 million.

#### Cell Line Flow cytometry

For surface MHC-II staining, cells were stimulated with recombinant human IFN-γ (10 ng/mL, 72 h) or vehicle control as described above. Cells were harvested by trypsinization, washed with PBS containing 2% FBS, and incubated with DAPI (1:10,000 dilution) and anti–HLA-DR antibody (Monoclonal Mouse, Anti-Human, HLA-DP, DQ, DR Antigen, Clone CR3/43, Catalog# M0775) for 1 h on ice.

Cells were washed, resuspended in PBS-2% FBS, and analyzed on a BD LSR-Fortessa instrument. Compensation and gating were performed in FlowJo software (v10.8.1); single, live (DAPI-negative) cells were gated for surface HLA-DR analysis.

## Computational methods

### Bulk sequencing data processing

#### RNA-seq analysis

Read alignment, quality control, and primary analysis were performed using the Visualization Pipeline for RNA-seq (VIPER)(47). Reads were aligned to hg38 using STAR v2.7.0f(48) followed by transcript assembly with cufflinks v2.2.1(49) and quality assessment with RSeQC v2.6.2. Differential gene expression was performed on raw read counts using DESeq2 v1.18.1(50) . GSEA was performed using the Broad GSEA Java application v4.1.0(51) with the MSigDB Hallmark collection.

#### ATAC-seq analysis

All samples were processed through the CHIPS computational pipeline developed at the Dana-Farber Cancer Institute Center for Functional Cancer Epigenetics. Sequence tags were aligned with Burrows-Wheeler Aligner (BWA) to hg38 and uniquely mapped, non-redundant reads were retained. Peak calling was performed with MACS2 v2.1.1.20160309, narrow peak option, q-value (FDR) threshold of 0.0175. BigWig tracks were generated with the bigwig average function in deeptools v3.5.1 and visualized in IGV v2.14.1. For differential accessibility analysis, peaks from all samples were processed as follows in our COBRA pipeline(52): peaks were merged to create a union peak set, sequencing depth normalized per sample, and DESeq2 used to determine differential peaks. Log fold changes were shrunk with lfcShrink for more accurate effect-size estimation. Motif analysis on differential peaks was performed with findMotifsGenome.pl in HOMER v3.0.0 at q ≤ 1 × 10⁻¹⁰. Sample signal at differential sites was visualized with deeptools. Principal component analysis used all peaks in the union set.

### Single-cell genomics analysis

#### snRNA-seq and scRNA-seq data processing

Sequencing reads were aligned to the GRCh38 reference using Cell Ranger version 7.1.0 (10x Genomics). For FLEX libraries (internal NMIBC and MIBC snRNA-seq), the multi-sample probe-aware pipeline was used with the 10x prebuilt FLEX human probe set reference. For standard 3′/5′ libraries (external MIBC and TME-enriched cohorts), the prebuilt hg38 10x reference was used. Doublets were identified and removed using scDblFinder v1.18.0 with default parameters. Nuclei or cells passing the following thresholds were retained: 500<UMI<80000; genes> 350; MT reads<2%. For the external cohort, we downloaded their processed and annotated rds file, without further filtering resulting in 67,988 cells.

#### Dimensionality reduction, integration, and clustering

Count matrices were normalized and variance-stabilized using Seurat v5.5.0 with SCTransform. Samples were merged and the top 25 principal components were used for nearest-neighbor graph construction (FindNeighbors) and UMAP embedding. Clusters were identified with the Louvain algorithm (FindClusters) at resolution 0.4.

#### Cell type annotation

Major lineage assignment (epithelial, T cell, B cell, macrophage, fibroblast, endothelial) was performed by manual inspection of canonical marker gene expression: EPCAM, KRT5, KRT20, UPK1B (epithelial); CD3D, CD3E (T cells); MS4A1, CD79A (B cells); CD68, CD163, C1QA (macrophages); COL1A1, DCN (fibroblasts); VWF, PECAM1 (endothelial). InferCNV (https://github.com/broadinstitute/inferCNV) (v1.19.1) was used to identify CNA in single cells with threshold: cutoff = 0.1, HMM = TRUE, leiden_method=“simple,” cluster_by_groups=TRUE, denoise=TRUE. Cells with extensive aneuploidy consistent with copy-number alterations were classified as malignant epithelial cells; cells with diploid profiles were classified as non-malignant urothelium.

#### T-cell subclustering and annotation

In the TME-enriched cohort, T cells (CD3D+, CD3E+) were re-extracted and re-clustered as above. Subtypes were assigned by canonical markers: CD8 Effector (CD8A, GZMB, PRF1, GZMH), CD8 Exhausted (CD8A, PDCD1, TOX, TIGIT), CD8 Mem-like (CD8A, IL7R, TCF7, SELL), CD4 Naïve-like (CD4, CCR7, TCF7, LEF1), CD4 Treg (CD4, FOXP3, IKZF2), CD4 Exhausted (CD4, PDCD1, CXCL13), Cycling T (MKI67, TOP2A), gd T (TRDC, TRGC1), and Thsp (HSPA1A, HSPA1B, HSPA6, IFNG). The Thsp subset was defined by co-expression of IFNG and heat-shock chaperones without canonical exhaustion markers (PDCD1, TOX).

#### Single-cell differential expression

Differential expression between malignant epithelial cells from tsMHC-II–high and tsMHC-II–low tumors was performed at the single-cell level using Seurat FindMarkers/FindAllMarkers with the Wilcoxon rank-sum test. Sample-level tsMHC-II status was assigned based on matched bulk RNA-seq GSVA scores and bimodal classification (see “MHC-II GSVA scoring and bimodal classification”). To validate results at the sample level and account for inter-sample variability, pseudobulk expression profiles were generated by aggregating raw counts of malignant epithelial cells per sample using Seurat AggregateExpression, followed by differential expression analysis using DESeq2 (v1.18.1). The data was then transferred to Scanpy using the zellkonverter package for final plots.

#### Paired scRNA-seq/scTCR-seq analysis

Feature matrices from CellRanger (version 7.1.0) were read in with Scanpy (version 1.15.1) and concatenated together. Cells were filtered based on high levels of mitochondrial counts and extreme total read counts (very high or low). Genes were filtered to those with at least 50 total counts, resulting in a final dataset of 84,051 cells and 19,413 genes. Expression counts were normalized using standard library size normalization, and then additionally log normalized, and z-scored. PCA was run on the z-scored, log-library size normalized counts. Harmony batch effect correction was run to align 2 separate sequencing runs (one with 4 samples, the other with 10. The kNN graph was generated using 100 nearest neighbors, 50 PCs, cosine metric on the batch-corrected PCA matrix, UMAP and Leiden clustering were run using this neighborhood graph.

The TCR data was read in using the scirpy package (v0.21.0) to confirm that the 8 clusters with high levels of expression of T cell marker genes also contained higher fractions of TCR sequences identified. T cells were selected cells for further analysis using these 8 clusters, and selecting only cells with TCR identified, resulting in a T cell dataset of 27,920 cells.

A new neighborhood graph was generated on this smaller T cell subset, sugin 60 neighbors, 50 PCs, cosine metric on the batch-corrected PCA matrix. Leiden clustering was run using this kNN graph and a new UMAP was generated. T cell subsets were annotated based on marker gene expression. A subset of clusters were difficult to annotate, as they seemed to express CD4, CD8A, and CD8B. These 7,539 cells were further subsetted, and a new neighborhood graph built using 60 nearest neighbors, 50 PCs, the Harmony batch-corrected matrix, cosine metric. Leiden clustering with resolution 1 and UMAP were then run, which enabled annotations of these cells into separate CD4 and CD8 populations based on marker gene expression.

TCR analysis was performed using the scirpy package (v0.21.0). Standard processing was run: the filtered_contig_annotations matrices were read in and concatenated together, chains were indexed and QCd. Clonotypes were determined based on the nucleotide sequences. Expanding clones were defined as those with a larger number of counts post-treatment than pre-treatment in one patient but were identified at both timepoints. Thsp clones were defined as those containing at least one Thsp T cell at either time point.

To determine the significance of sharing of CD4 vs CD8 T cells with these Thsp clones, a hypergeometric test was run for each patient in a cell-based manner. We tested for enrichment of either CD4 cells (from the CD4 Treg, CD4 Naïve-like, CD4 Exhausted populations) or CD8 cells (from the CD8 Mem-like, CD8 Effector, CD8 Exhausted populations) within cells clonally shared with Thsp cells (i.e., cells part of Thsp-containing clones that were composed of strictly more than 1 cell). The results for CD4 populations were not significant in any patient, with FDR=1, but the results for CD8 populations were significant in all patients with FDR < 1.3e-54.

To determine the significance of sharing of CD4 vs CD8 T cells with these Thsp clones across the 8 pre-treatment samples, we first used a 1-sided paired Wilcoxon/T Test to test that the fraction of CD4 T cells (from the CD4 Treg, CD4 Naïve-like, CD4 Exhausted populations) that are part of a clone (with 2+ cells) is significantly higher than the fraction of CD4 T cells that are part of a Thsp clone (with 2+ cells), FDR < 0.025/0.017(Wilcoxon/T respectively). In contrast, we used a 1-sided paired Wilcoxon/T Test to test that the fraction of CD8 T cells (from the CD8 Mem-like, CD8 Effector, CD8 Exhausted populations) that are part of a Thsp clone (with 2+ cells) clone is significantly higher than the fraction of CD8 T cells that are part of a (with 2+ cells)., FDR < 0.025/0.015(Wilcoxon/T respectively).

#### Chu CD8 atlas projection

CD8-lineage T cells from the TME-enriched cohort were projected onto the Chu et al. pan-cancer CD8 reference(33) using their TCM approach to align and map T cells in a scRNA-seq dataset to their T cell atlas with some small adjustments to their code to make it run. Each query cell received a predicted Chu CD8 subset label and probability score. For the cross-cohort Tstr abundance comparison (Figure 3i), bladder cancer was added as a 16th cancer type alongside the 15 cancer types in the Chu atlas, and the mean fraction of CD8 cells annotated as Tstr per tumor type was computed.

#### scATAC-seq data processing

The two internal NMIBC scATAC-seq samples were processed using the cellranger-atac (v2.0.0) pipeline with default parameters. Quality control filtering of low-quality cells was performed using the R packages Seurat (v3) and Signac (v1.6.0) based on criteria including nucleosome-binding pattern strength, transcription start site enrichment score, number of fragments in peaks>100, and peaks > 600. For the nine MIBC scATAC-seq samples from Sundaram et al., the scATAnno approach(53) was used to perform cell type annotation. For both datasets, accessibility at MHC-II loci (CD74, CIITA, HLA-DRA, HLA-DRB1, HLA-DQB1, HLA-DPA1) was visualized in IGV v2.14.1 using sample-aggregated pseudobulk tracks restricted to tumor cells.

### Cohort-level computational analyses

*MHC-II GSVA scoring and bimodal classification.* For each bulk RNA-seq cohort (UROMOL, TCGA, PURE-01, Bellmunt internal), expression matrices were log-transformed and Gene Set Variation Analysis (GSVA; R package GSVA v2.4.9)(22) was applied using a comprehensively curated 17-gene MHC-II signature. Rather than relying on a generalized pathway definition, this specific panel was manually curated from standard reference molecular databases (including the Reactome MHC Class II Antigen Presentation pathway) to isolate the core functional machinery of human MHC-II restricted antigen processing. The 17-gene panel explicitly encompasses the classical HLA heterodimers (HLA-DRA, HLA-DRB1, HLA-DRB3, HLA-DRB4, HLA-DRB5, HLA-DPA1, HLA-DPB1, HLA-DQA1, HLA-DQA2, HLA-DQB1, HLA-DQB2), the essential intracellular accessory and loading molecules (HLA-DMA, HLA-DMB, HLA-DOA, HLA-DOB, CD74), and the major histocompatibility complex transactivator master regulator (CIITA). Score distributions were modeled as a mixture of two Gaussian components using mixdist v0.5.5, and the intersection between the two component densities (computed with rootSolve v1.8.2.4) was used as a data-driven cutpoint to assign MHC-II–high versus MHC-II–low status. The same procedure was applied at the single-cell level by computing GSVA scores per sample on pseudobulked counts for sample-level classification.

#### Whole-genome doubling annotation

WGD status for UROMOL samples was obtained from the published multi-omic source data accompanying the comprehensive genomic characterization of the UROMOL cohort(36). Specifically, WGD classifications were extracted directly from the study’s curated source data files covering Figures 1–5 and Extended Data Figures 1–10. Samples without a confident WGD call were excluded from the WGD–MHC-II analysis.

#### Consensus in vitro IFN-γ response signature

For each of the six bladder cancer cell lines (J82, TCCSUP, SCaBER, 5637, RT4, SW780), differential expression between IFN-γ-stimulated (10 ng/mL, 72 h; 48 h for RT4) and unstimulated conditions was computed with DESeq2 v1.18.1. The five lines showing coordinated MHC-II induction (J82, TCCSUP, SCaBER, 5637, RT4) were designated as responsive; SW780 was excluded from signature derivation. The consensus IFN-γ UP signature was defined as genes meeting the following criteria: (i) median log₂ fold change > 1 across the five responsive lines, (ii) significantly upregulated (DESeq2 adjusted p < 0.05) in at least 5 lines, and (iii) combined p-value < 0.05 using Fisher’s method. The IFN-γ DOWN signature was defined analogously with the opposite fold-change direction. The final UP and DOWN gene lists are provided in Supplementary Table 2.

#### In vitro to in vivo concordance analysis

For Figure 2e, the consensus in vitro IFN-γ UP signature was tested for enrichment among genes upregulated in MHC-II–positive versus MHC-II–negative malignant epithelial cells in the internal snRNA-seq cohort using the fgsea R package with 10,000 permutations. Genes were ranked by log2FC.

The in vitro DOWN signature was tested analogously. For the genome-wide concordance analysis (Figure 2f), gene-level log₂ fold changes from the in vitro DE analysis (median across five responsive lines) and the in vivo DE analysis (MHC-II– positive vs. negative malignant cells, p < 0.05) were intersected, and Spearman’s rank correlation was computed across all genes detected in both settings.

#### tsMHC-II program signature construction

The tsMHC-II program signature was constructed as the three-way intersection of gene sets defined within this study, each comprising genes significantly upregulated in a distinct comparison: (i) IFN-γ-stimulated versus unstimulated bladder cancer cell lines, requiring significance in at least 5 of 6 lines tested (J82, TCCSUP, SCaBER, 5637, RT4, SW780; 10 ng/mL IFN-γ for 72 h; 48 h for RT4) at DESeq2 v1.18.1 adjusted p < 0.05 with no fold-change threshold; (ii) tsMHC-II–positive versus tsMHC-II–negative malignant epithelial cells in the internal NMIBC snRNA-seq cohort, at adjusted p < 0.05 (Seurat FindMarkers, Wilcoxon rank-sum test on log-normalized counts), with no fold-change threshold; and (iii) the same comparison in the external MIBC snRNA-seq cohort using the same DE method and thresholds. Sample-level tsMHC-II status for the snRNA-seq comparisons was assigned by bimodal Gaussian classification of the matched bulk RNA-seq MHC-II GSVA score. The three-way intersection yielded 11 genes: CD74, CIITA, CXCL10, CXCL11, ICAM1, GBP5, IL32, SOD2, HAPLN3, TNFRSF1B, and SLC12A7. The full input gene lists and the final 11-gene signature are provided as Supplementary Tables 3-6.

#### tsMHC-II program scoring and stratification

The tsMHC-II program score was computed by applying GSVA (R package GSVA v2.4.9) to log-transformed bulk RNA-seq expression matrices from the PURE-01 cohort using the curated 11-gene signature.

Score distributions in each cohort were modeled as a mixture of two Gaussian components using mixdist v0.5.5, and the intersection between component densities (computed with rootSolve v1.8.2.4) was used as a data-driven cutpoint to assign MHC-II–high and MHC-II–low status.

### Survival, statistical, and clinical analyses

#### Survival analyses

Kaplan-Meier curves were generated with the survival v3.7.0 and survminer v0.4.9 R packages and compared using the log-rank test. Hazard ratios and 95% confidence intervals were estimated by Cox proportional-hazards regression. The proportional-hazards assumption was assessed by testing for non-zero slopes in the Schoenfeld residuals over time using the cox.zph function, and the assumption was met across all evaluated baseline multivariate models. Univariate Cox models were fit separately for each predictor. Multivariate Cox models were specified as follows:

- UROMOL (luminal (216) and basal(218) stratification): PFS ∼ MHC2_status (MHC2- vs MHC2+ reference group). Subtype-specific hazard ratios and 95% confidence intervals were extracted using the broom package, and a stratified log-scale forest plot framework was engineered using ggplot2 to display the consolidated estimates across differentiation states without reference rows.
- PURE-01 (n = 82): RFS ∼ tsMHC-II program status + TMB (binarized at published cutoff at 10) + PD-L1 expression categorical at CPS ≥ 10.

A formal subtype × MHC-II interaction term was tested in UROMOL via Cox regression on PFS.

#### Tumor purity sensitivity analysis (PURE-01)

Tumor purity estimates were obtained from the FoundationOne (FMOne) sequencing of pre-treatment specimens. To test whether the bulk tsMHC-II program signal was confounded by tumor purity, the program GSVA score was regressed on tumor purity by linear regression and the Pearson correlation between score and purity was computed. The residuals from this regression were retained as a purity-independent measure of program activity, and tumors were classified as program-high (residual ≥ 0; n = 40) or program-low (residual < 0; n = 42). Recurrence-free survival was compared between these groups by the log-rank test and Cox proportional-hazards regression, both unadjusted and in a multivariate model including TMB (binarized at 10 mut/Mb) and PD-L1 (CPS ≥ 10) as specified above. The proportional-hazards assumption was assessed using cox.zph.

#### IMvigor210 analysis

Processed expression and clinical data were obtained from the IMvigor210CoreBiologies package. Analyses were restricted to patients with pretreatment expression data, overall survival data, and a profiled specimen from the primary tumor site. Specimens from lymph node and metastatic sites were excluded on measurement-validity grounds, as bulk MHC-II signal in these tissues derives substantially from constitutively MHC-II-positive resident immune populations. Tissue of origin was used to define both the exclusion and the bladder versus upper tract classification. tsMHC-II program GSVA scores were computed on the resulting 288- sample matrix using the same 11-gene signature applied to PURE-01, and a two-component Gaussian mixture was fitted to the 288 scores with the intersection of the component densities taken as the cutpoint (0.007). Only one of 288 tumors was classified differently under this cutpoint than under a mixture fitted to all 348 samples.

Overall survival was modeled with Cox proportional hazards regression, unadjusted and adjusted for ECOG performance status, liver metastasis, prior platinum therapy, and sex. Hemoglobin is not available in this dataset, so the complete Bellmunt risk score could not be computed; adjustment is therefore described as adjustment for available baseline prognostic factors. Site dependence was tested with a site-by-program interaction term and evaluated by likelihood ratio test. Proportional hazards assumptions were assessed with scaled Schoenfeld residuals and were satisfied in all models (global p > 0.19). All models were fitted with both the binary classification and the continuous standardized score.

#### Other statistical analyses

Categorical associations between patient subsets, therapeutic response groups, and clinical stages were assessed by Fisher’s exact test or the chi-square test. Continuous comparisons of signature scores between two groups (such as objective responders versus non-responders) were performed using the Wilcoxon rank-sum test. Bar charts tracking response distribution percentages were constructed using ggplot2. Correlations were computed with Pearson’s coefficient when both variables were approximately normal and Spearman’s coefficient otherwise. All p-values are two-sided unless otherwise stated, and significance was defined as p < 0.05. Analyses were performed in R v4.3.2.

## Data availability

Raw and processed sequencing data generated in this study (bulk RNA-seq of bladder cancer cell lines, ATAC-seq of bladder cancer cell lines, snRNA-seq of NMIBC and MIBC tumors, scRNA-seq and scTCR-seq of TME-enriched bladder tumors) have been deposited at GSE334331. Public datasets reanalyzed in this study are detailed in the methods section.

## Code availability

The VIPER RNA-seq pipeline is available at https://bitbucket.org/cfce/viper/src/master/; the CHIPS ATAC-seq pipeline is available at https://github.com/liulab-dfci/CHIPS. All other analyses used published R packages and Bioconductor tools at the versions specified in the relevant subsections above.

## Acknowledgements

H.W.L. acknowledges funding from The Massachusetts Life Sciences Corporation under the Bits to Bytes program. P.C. acknowledges funding from the Ministry of Economy and Competitiveness, Institute deSalud Carlos III (Institute of Health Carlos III)— PI23/01533. J.B. acknowledges the support from the Kaifer Family Bladder Research Fund. C.C. acknowledges support from Fundació La Marató de TV3 through grant 201908-30-31, which supported this research.

