## Supplementary Figures for "A tumor-cell MHC-II program is associated with checkpoint-blockade outcomes across stages of bladder cancer"

### Supplementary Figure 1

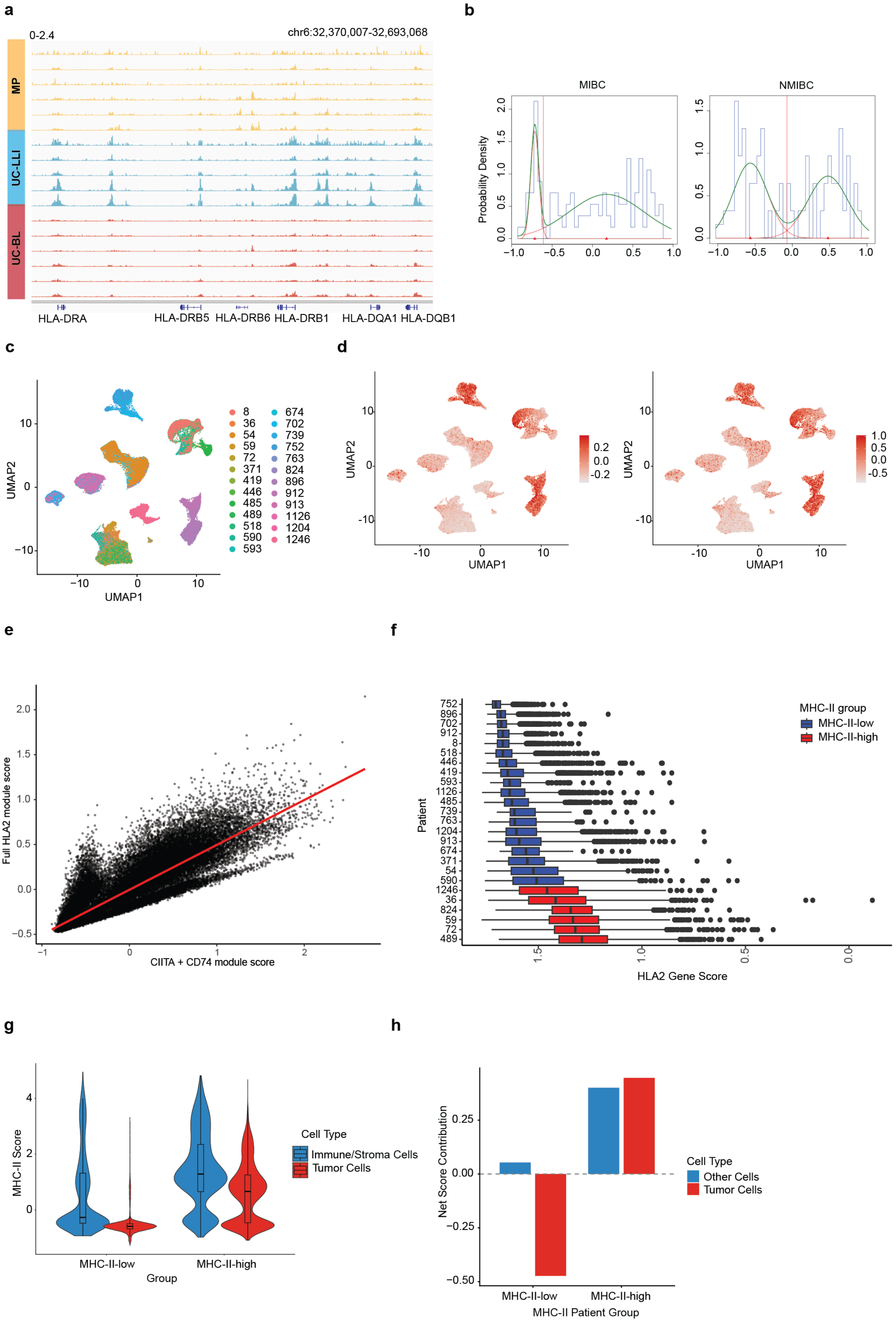

**Supplementary Figure 1. MHC-II pathway bimodal distribution and validation in an independent MIBC scRNA-seq cohort.**

**(a)** IGV browser tracks showing H3K27ac ChIP-seq signal at MHC-II gene loci on chromosome 6 across three NMIBC molecular subtypes: micropapillary (MP), urothelial carcinoma–basal-like (UC-BL), and urothelial carcinoma–luminal-like with immune infiltration (UC-LLI). H3K27ac enrichment at MHC-II promoters is highest in the UC-LLI subtype, indicating active chromatin marks at these loci specifically in luminal tumors with immune features. Data are from FiTAc-seq profiling of 16 NMIBC FFPE specimens (21).

**(b)** Distribution of GSVA-derived MHC-II pathway signature scores in the PURE-01 MIBC cohort (n = 82; left) and the Bellmunt NMIBC cohort (n = 62; right). Histograms show original score distributions. Green curves indicate fitted bimodal Gaussian mixture models; red vertical lines mark the inter-peak minima used as data-driven cutpoints for MHC-II–high/low classification (see Methods). Both cohorts recapitulate the bimodal distribution observed in the UROMOL and TCGA datasets (Figure 1a).

**(c)** UMAP visualization of 67,988 cells from an independently published MIBC scRNA-seq cohort (n = 25 patients, 10x 3′ chemistry), showing tumor cells colored by patient identity. Data were processed through the same analytical pipeline as the internal NMIBC cohort (Methods). Comparable clustering and cell-type distributions confirm cross-platform consistency.

**(d)** UMAP visualization of tumor cells in the external MIBC scRNA-seq cohort. Left: cells colored by aggregate expression of all MHC-II pathway–associated genes. Right: cells colored by *CIITA* and *CD74* expression only. Multiple patients show extensive tumor-intrinsic MHC-II expression, consistent with the NMIBC observations.

**(e)** The CD74 + CIITA proxy score closely tracks the full MHC-II pathway signature. Per-cell correlation between the two-gene proxy score (CD74 + CIITA module score, x-axis) and the full MHC-II pathway module score (y-axis) in the external 25-case MIBC scRNA-seq cohort, which provides complete coverage of the MHC-II gene set. Each point represents a single cell; the red line shows the linear fit (Pearson $r = 0.90$, $p < 0.001$). The strong correlation confirms that the two-gene proxy used in the FLEX-based internal cohort faithfully captures coordinated MHC-II pathway activity.

**(f)** Box plot showing per-patient MHC-II pathway gene expression scores across samples in the external MIBC cohort. MHC-II–high samples (red) and MHC-II–low samples (blue) were classified using the same bimodal thresholding approach as in the NMIBC cohorts.

**(g)** Per-cell MHC-II score by compartment and MHC-II group. Violin plots of the MHC-II proxy score (CD74 + CIITA module score) in immune/stromal cells (blue) and tumor cells (red), separated by MHC-II–low and MHC-II–high tumor groups in the internal snRNA-seq cohort. Boxplots indicate median and interquartile range. Immune and stromal cells display the highest per-cell scores in both groups, while tumor cells shift from a predominantly negative distribution in MHC-II–low tumors to a positive distribution in MHC-II–high tumors. Group means, medians, and cell counts: MHC-II–low immune/stroma, mean 0.588, median −0.270, n = 1,827; MHC-II–low tumor, mean −0.520, median −0.587, n = 18,301; MHC-II–high immune/stroma, mean 1.43, median 1.28, n = 1,888; MHC-II–high tumor, mean 0.621, median 0.664, n = 4,835.

**(h)** Net contribution of each compartment to the tissue MHC-II signal. Net contribution calculated as the product of compartment proportion and mean per-cell MHC-II score, shown separately for MHC-II–low and MHC-II–high tumors. In MHC-II–low tumors, tumor cells contribute negatively to the aggregate signal; in MHC-II–high tumors, they become the dominant positive contributor. The immune/stromal contribution changes comparatively little between groups, indicating that the between-group difference in MHC-II signal is driven primarily by the shift in tumor-cell expression.

#### Supplementary figure 2

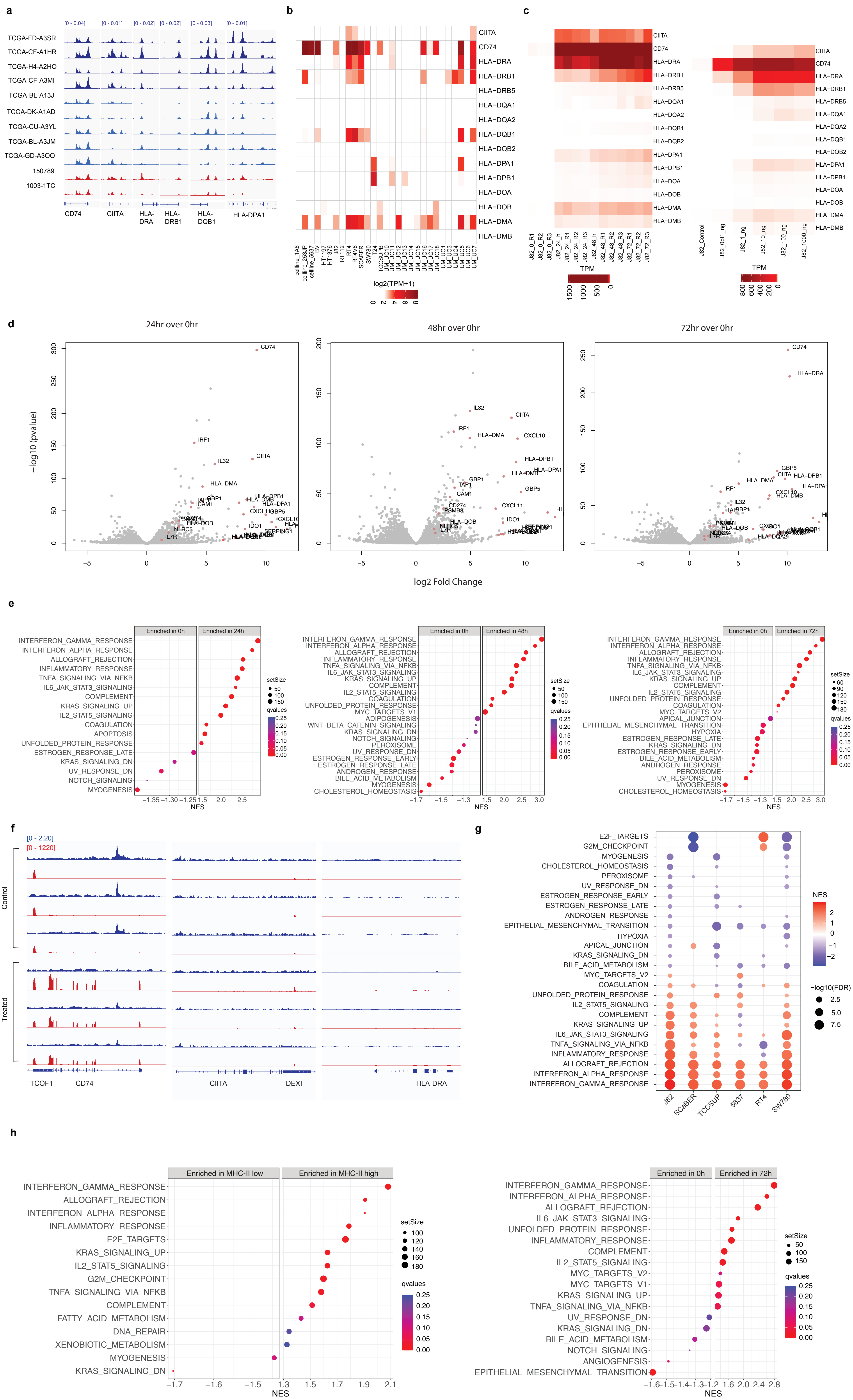

#### SUPPLEMENTARY Figure 2

i

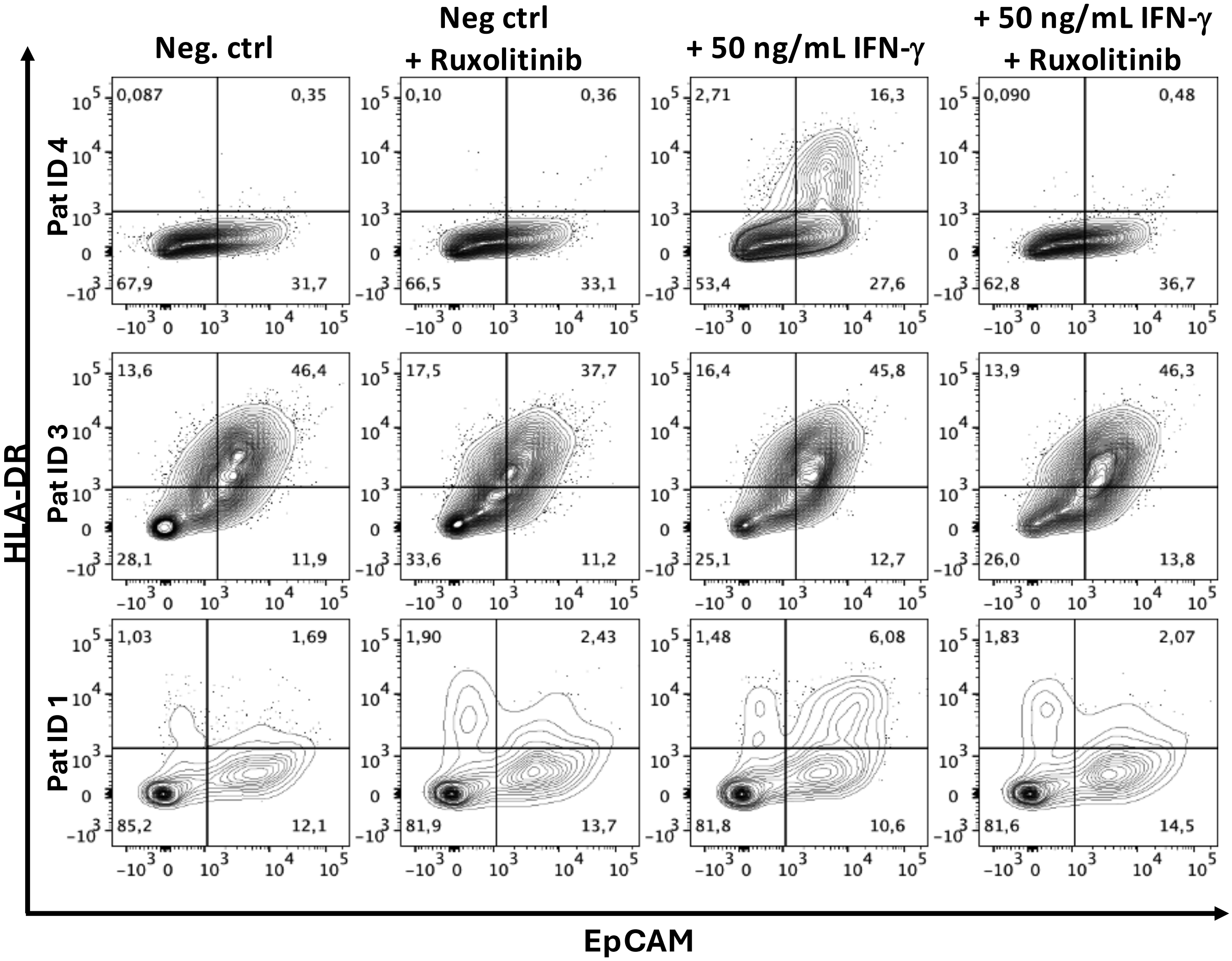

**Supplementary Figure 2. IFN-γ dose–response, time-course, and pathway analysis in bladder cancer cell lines.**

**(a)** Chromatin accessibility at MHC-II loci in primary bladder tumors. IGV browser tracks showing tumor-cell scATAC-seq accessibility at MHC-II loci (CIITA, CD74, HLA-DRA, HLA-DRB1, HLA-DQB1, HLA-DPA1) in nine published MIBC samples (Sundaram et al. 2024; upper, blue) and two internal NMIBC samples (lower, red). Tracks are sample-aggregated pseudobulk signal restricted to tumor cells. Most samples show accessible chromatin at MHC-II promoters across both disease stages, mirroring the cell-line data. The MIBC samples are ordered by GSVA-derived MHC-II enrichment from Figure 1a, from most to least enriched. MHC-II-negative samples show lower accessibility, though the difference does not reach statistical significance.

**(b)** Heatmap of RNA-seq expression values (TPM) for MHC-II pathway genes across 30 bladder cancer cell lines from the publicly available dataset GSE97768. Rows represent genes; columns represent cell lines. Expression of canonical MHC-II genes (HLA-DRA, HLA-DRB1, CIITA) is negligible across the panel, confirming that constitutive MHC-II transcription is rare in bladder cancer cell lines. Limited CD74 expression is observed in a subset of lines but does not translate into coordinated MHC-II complex expression.

**(c)** Left: heatmap of RNA-seq expression values (TPM) for MHC-II pathway genes in J82 cells across increasing IFN-γ doses (0, 0.1, 1, 10, 100, and 1000 ng/mL) at 72 h. IFN-γ elicits dose-dependent transcriptional activation of MHC-II genes, reaching a plateau at approximately 10 ng/mL. Right: heatmap of RNA-seq expression values (TPM) for MHC-II pathway genes in J82 cells across IFN-γ stimulation time points (0, 24, 48, and 72 h at 10 ng/mL). Rows represent individual genes; columns represent time points. Color intensity reflects expression magnitude. Coordinated transcriptional activation of MHC-II pathway genes is evident by 24 h and sustained through 72 h.

**(d)** Volcano plots of differentially expressed genes in J82 cells at 24 h, 48 h, and 72 h relative to 0 h (untreated). The x-axis shows $\log_2$ fold change; the y-axis shows $-\log_{10}$ adjusted p-value. Significantly upregulated genes are shown in red; downregulated genes in blue (thresholds: FDR < 0.05, $|\log_2 FC| > 1$). The number of differentially expressed genes increases with stimulation duration.

**(e)** GSEA results showing significantly enriched and depleted MSigDB Hallmark gene sets in J82 cells at 24 h, 48 h, and 72 h versus 0 h. Gene sets are ranked by normalized enrichment score. IFN-γ response pathways show progressive enrichment across time points.

**(f)** IGV browser tracks showing ATAC-seq chromatin accessibility (blue) and RNA-seq signal (red) at CIITA, HLA-DRA, and CD74 in SW-780 cells in control versus IFN-γ stimulation. All ATAC-seq tracks share a common y-axis scale; all RNA-seq tracks share a separate scale (values indicated in the upper-left corner). IFN-γ does not induce de novo chromatin opening at MHC-II promoters in SW-780. CD74 shows some transcriptional increase, but induction of CIITA and HLA-DRA is minimal.

**(g)** GSEA results showing significantly enriched and depleted MSigDB Hallmark gene sets in the comparison between IFN-γ-stimulated and control cells across all six cell lines.

**(h)** GSEA results showing significantly enriched and depleted MSigDB Hallmark gene sets in (left) the comparison between MHC-II-high and MHC-II-low malignant epithelial cells from the internal snRNA-seq cohort and (right) the average in vitro response used in the correlation plot in Figure 2f. This analysis identifies pathway-level transcriptional concordances between the MHC-II states of patient tumor cells and IFN-γ-stimulated cell lines.

**(i)** Ex vivo validation of tumor-intrinsic MHC-II expression in primary bladder tumors. Fresh tumor specimens from three patients were dissociated and cultured ex vivo. Twenty-four hours after isolation, cells were stimulated with recombinant human IFN-γ (50 ng/mL) for 72 h in the presence or absence of the JAK1/2 inhibitor ruxolitinib (20 μM). HLA-DR expression was assessed by flow cytometry within the viable $CD45^{-}EpCAM^{+}$ tumor-cell compartment. One tumor (Pat 3) displayed a pre-existing MHC-II-high phenotype, whereas the remaining two were MHC-II-low. IFN-γ induced HLA-DR upregulation in MHC-II-low tumors, while ruxolitinib completely abrogated this effect, confirming that tumor-cell MHC-II expression is regulated through canonical IFN-γ–JAK/STAT signaling.

#### Supplementary figure 3

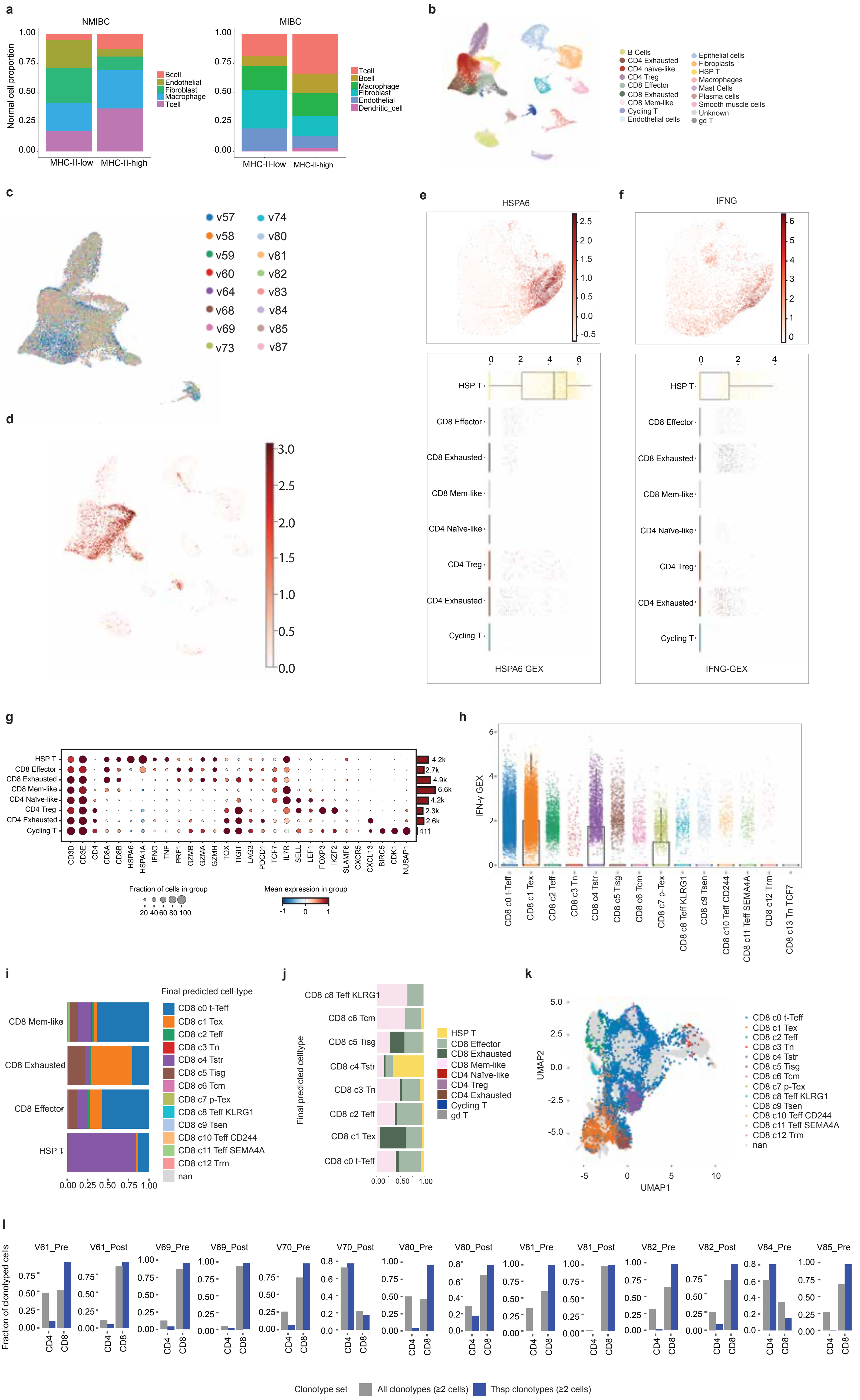

**Supplementary Figure 3. TME-enriched cohort characterization and Thsp population analysis.**

**(a)** Stacked bar plots showing the proportional composition of non-malignant immune and stromal cell subtypes in MHC-II–low versus MHC-II–high tumors, in the internal NMIBC snRNA-seq cohort (left) and the external MIBC cohort (Gouin et al. 2021; right). Malignant epithelial cells were excluded so that the comparison reflects microenvironment composition alone. In both cohorts, MHC-II–high tumors show increased T-cell and B-cell fractions and reduced fibroblast content, indicating a lymphocyte-enriched, less desmoplastic microenvironment. The overall distribution of non-malignant cell types differed significantly between MHC-II groups in each cohort (chi-square test: NMIBC $p < 2.2 \times 10^{-16}$; MIBC $\chi^2 = 686.1$, df = 5, $p < 2.2 \times 10^{-16}$). Cell-type-specific testing (Fisher's exact test) confirmed significant T-cell enrichment (NMIBC $p = 2.9 \times 10^{-41}$; MIBC $p = 5.1 \times 10^{-56}$), B-cell enrichment (NMIBC $p = 6.8 \times 10^{-16}$; MIBC $p = 3.0 \times 10^{-27}$), and fibroblast depletion (NMIBC $p = 4.8 \times 10^{-45}$; MIBC $p = 1.6 \times 10^{-62}$) in MHC-II–high tumors.

**(b)** UMAP visualization of 34,703 cells from the TME-enriched scRNA-seq cohort (n = 16 samples from fresh biopsies and cystectomy specimens), colored by annotated cell type. The cohort comprises >90% viable TME cells, providing substantially improved resolution of the immune compartment relative to the tumor-enriched snRNA-seq datasets.

**(c)** UMAP of all cells in the TME-enriched cohort, colored by patient identity. Multiple patients contribute to each annotated cell-type cluster, confirming that cell-type assignments are not driven by patient-specific batch effects.

**(d)** UMAP of all cells in the TME-enriched cohort, colored by IFNG expression. IFN-γ transcripts localize predominantly to the T-cell compartment.

**(e)** HSPA6 expression in the paired scRNA-seq/scTCR-seq cohort (eight MIBC patients, 14 samples with 6 pre- and post-NAC pairs). UMAP of T cells colored by HSPA6 (top) and HSPA6 expression across annotated T-cell subtypes (bottom). A discrete HSPA6-high population is present, mirroring the Thsp cluster of the larger TME-enriched cohort.

**(f)** IFNG expression in the same paired cohort. UMAP of T cells colored by IFNG (top) and IFNG expression across annotated T-cell subtypes (bottom). IFN-γ concentrates in the same population as HSPA6 (panel e), confirming that the IFN-γ–high, heat-shock–high Thsp population is recapitulated in this cohort.

**(g)** Dot plot of representative marker genes across annotated T-cell subtypes in the paired cohort. Dot color represents mean normalized expression; dot size represents the fraction of cells expressing the gene. The canonical CD8 effector, exhausted, memory-like, CD4, and Thsp subtypes are all recovered, confirming that the T-cell architecture recapitulates that of the larger TME-enriched cohort.

**(h)** Re-analysis of the Chu et al. pan-cancer CD8 T-cell atlas. IFNG expression across CD8 T-cell states shows that IFN-γ production is concentrated in the stress-response (Tstr) and exhausted (Tex) states, consistent with the IFN-γ–high Thsp population defined here.

**(i)** Proportional mapping of each of our CD8-lineage annotations (rows) onto Chu et al. CD8 atlas states (colors). Approximately 80% of Thsp (HSP T) cells are assigned to the Tstr state (CD8_c4_Tstr).

**(j)** Reverse mapping: Chu CD8 atlas states (rows) colored by our annotations. Approximately 65% of Chu Tstr (c4_Tstr) cells are annotated as Thsp.

**(k)** UMAP of our CD8-lineage T cells projected onto the Chu CD8 reference (refUMAP coordinates), colored by assigned Chu atlas state (Methods).

**(l)** Per-sample TCR clonotype lineage composition for each of the 14 samples from eight MIBC patients. For each sample, bars show the fraction of clonotyped cells annotated as CD4 or CD8 subsets across all clonotypes containing at least two cells (gray) and across clonotypes containing at least one Thsp cell (blue). Thsp cells are excluded from both numerators. Sample labels denote patient and specimen type, with paired patients contributing a pre-NAC transurethral resection sample and a post-NAC cystectomy sample. The enrichment of CD8 subsets in Thsp-containing clonotypes is consistent across 13 of 14 samples. Summary values are shown in Figure 3f.

### Supplementary figure 4

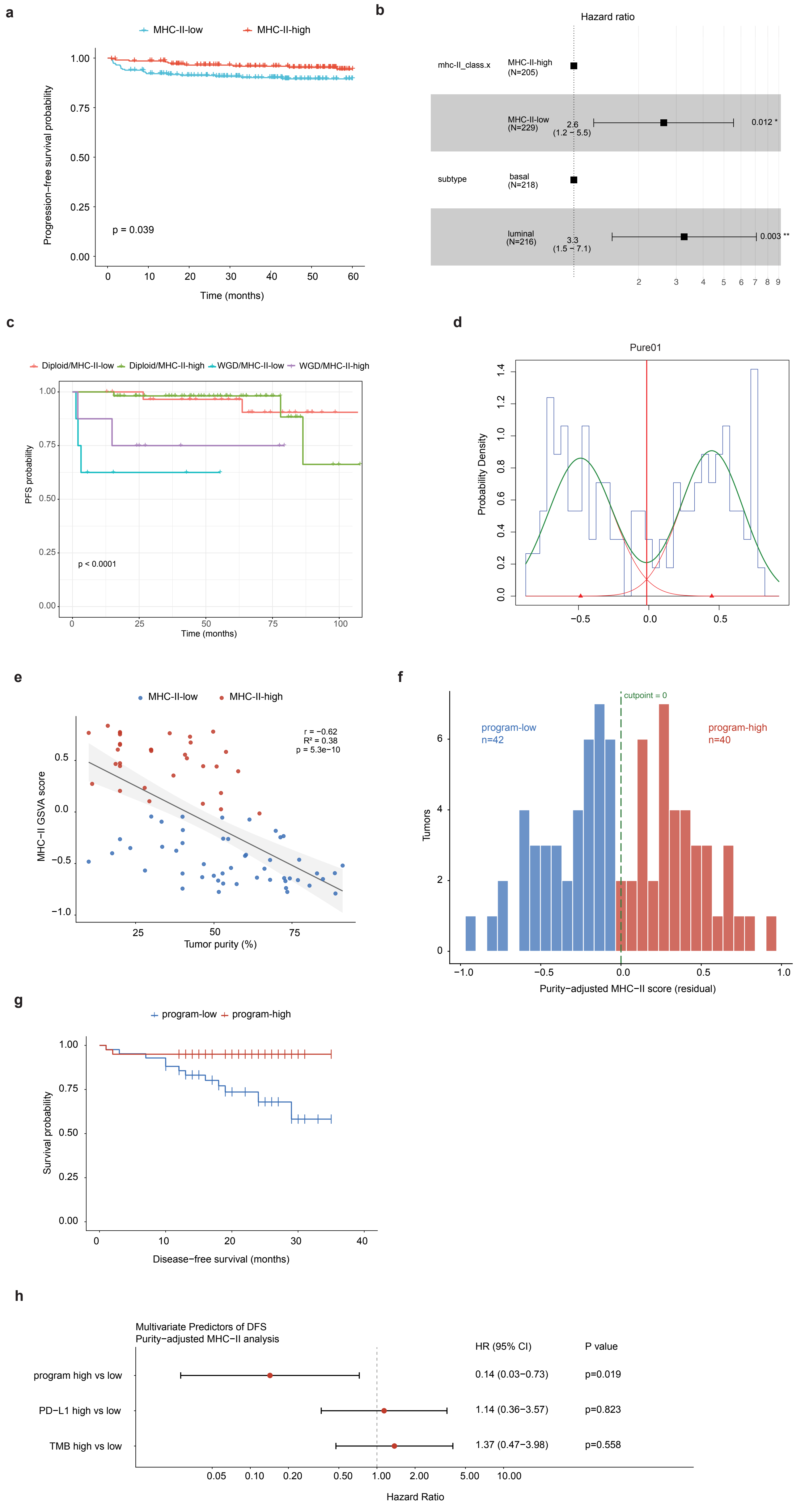

**Supplementary Figure 4. tsMHC-II expression is prognostic in NMIBC, associated with immunotherapy outcome in MIBC, and robust to tumor purity.**

**(a)** Kaplan-Meier curves for progression-free survival in the full UROMOL analytic cohort (n = 434) stratified solely by MHC-II status (MHC-II-high, n = 205; MHC-II-low, n = 229), without accounting for molecular subtype. Log-rank p = 0.039. Although MHC-II status reaches nominal significance in this unadjusted analysis, the effect is substantially more pronounced within the luminal subtype (Figure 4a, right), reflecting dilution of the signal by basal/MHC-II-low patients whose outcomes are comparable to both MHC-II-high groups.

**(b)** Forest plot from a Cox proportional hazards main effects model (n = 434, 33 progression events) showing the marginal association of MHC-II-low status with progression-free survival adjusted for molecular subtype (upper row), and the marginal association of luminal subtype adjusted for MHC-II status (lower row). MHC-II-high and basal subtype are the respective reference categories. MHC-II-low status is associated with a significantly higher risk of progression (HR 2.63, 95% CI 1.23 to 5.56, p = 0.012), and luminal subtype is associated with a significantly higher risk than basal subtype after adjustment for MHC-II status (HR 3.27, 95% CI 1.51 to 7.08, p = 0.003). Overall model likelihood ratio test p = 0.001. This model complements the subtype-stratified analysis in Figure 4b by showing the independent contributions of each feature to outcome in a single framework.

**(c)** Left: Kaplan-Meier curves for progression-free survival in luminal UROMOL tumors stratified jointly by MHC-II status and whole-genome doubling (WGD) status, yielding four groups: diploid/MHC-II-high (n = 54), diploid/MHC-II-low (n = 31), WGD/MHC-II-high (n = 8), and WGD/MHC-II-low (n = 8). Overall log-rank p < 0.0001. Both WGD-positive groups show worse progression-free survival than both diploid groups, consistent with the established adverse prognostic role of WGD in NMIBC. The WGD subgroups are small (n = 8 each) and findings within these groups should be interpreted cautiously.

**(d)** Distribution of GSVA-derived tsMHC-II program signature scores across all tumors in the PURE-01 MIBC cohort (n = 82). The histogram shows the observed score distribution; green curves show the fitted bimodal Gaussian mixture model; the red vertical line marks the inter-peak minimum used as a data-driven cutpoint to classify tumors as program-high or program-low (see Methods). The bimodal distribution indicates that tsMHC-II program activity segregates into two discrete transcriptional states in this cohort.

**(e)** Scatter plot of tsMHC-II program GSVA score (y-axis) against tumor purity (x-axis) in pre-treatment PURE-01 samples (n = 82). Each point is one tumor, colored by program status under the primary trough cutpoint (panel d; Figure 4). The solid line shows the linear regression fit with 95% confidence band. Program score is inversely correlated with purity (Pearson r = -0.62, $R^2 = 0.38$, $p < 0.001$), indicating that purity accounts for approximately 38% of inter-sample variance in the program score.

**(f)** Distribution of purity-residualized tsMHC-II program scores (residuals from the regression in panel e). Tumors are classified as program-high (residual ≥ 0; n = 40) or program-low (residual < 0; n = 42) about a cutpoint of zero (dashed line).

**(g)** Kaplan–Meier curves for recurrence-free survival in PURE-01, stratified by purity-residualized program status (program-high, n = 40; program-low, n = 42). The purity-independent component of the program score remains significantly associated with outcome (unadjusted HR 0.18, 95% CI 0.05 to 0.65, p = 0.009). The purity-adjusted estimate from the multivariate model in panel h is HR 0.14, 95% CI 0.03 to 0.73, p = 0.019.

**(h)** Forest plot of a purity-adjusted multivariate Cox model of recurrence-free survival in PURE-01 (n = 82, 14 events), including purity-residualized program status, PD-L1 (CPS ≥ 10), and TMB (≥ 10 mut/Mb). Program-high status is the only independent predictor of favorable outcome (HR 0.14, 95% CI 0.03 to 0.73, p = 0.019); PD-L1 (HR 1.14, 95% CI 0.36 to 3.57, p = 0.823) and TMB (HR 1.37, 95% CI 0.47 to 3.98, p = 0.558) are not significant. Hazard ratios are plotted on a log scale; points to the left of the dashed line favor program-high tumors.

### Supplementary figure 5

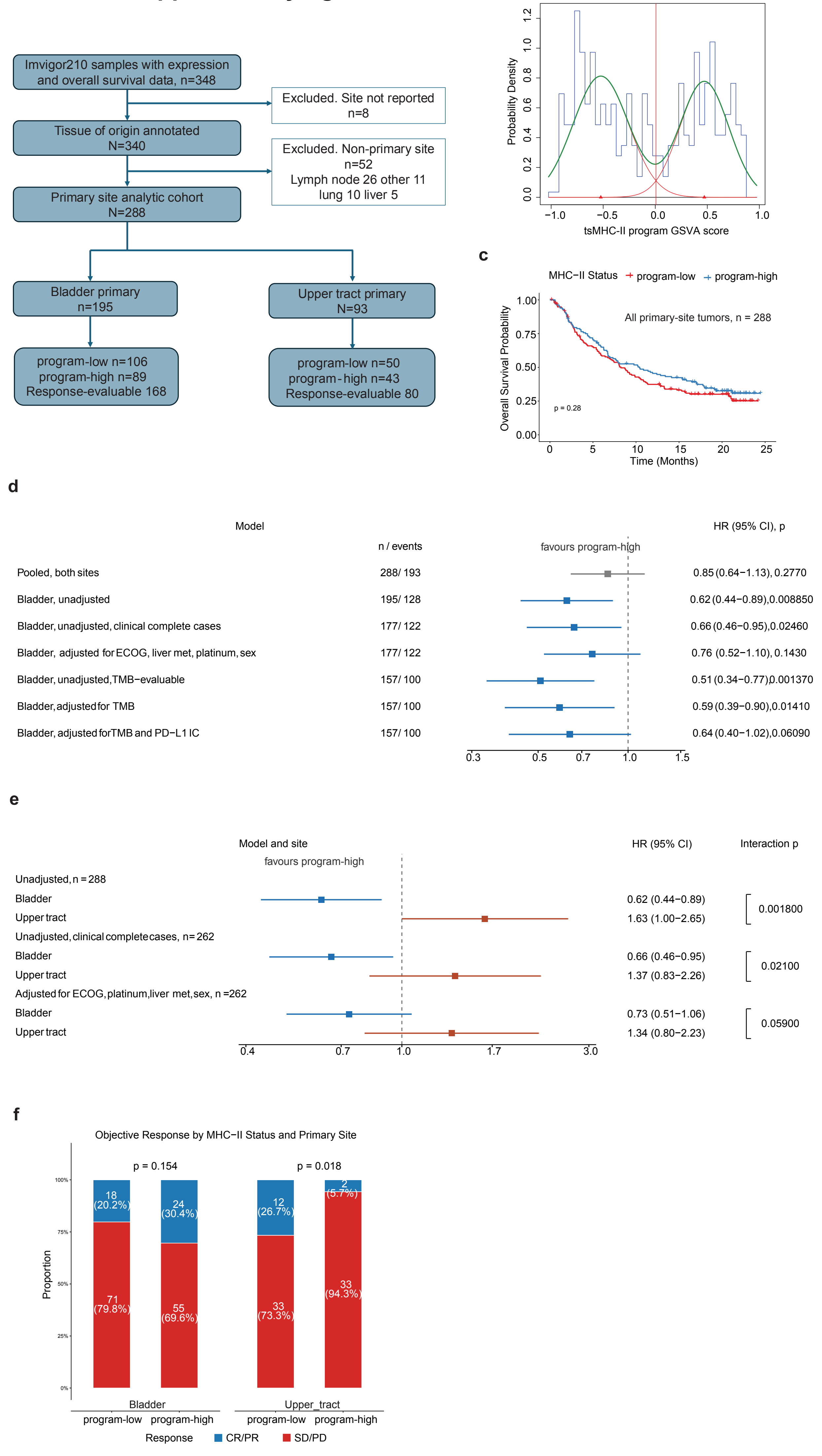

**Supplementary Figure 5. Cohort derivation, classification, and site-specific analyses in IMvigor210.**

**(a)** Derivation of the analytic cohort. Of 348 patients with pretreatment expression and overall survival data, 8 had no reported tissue of origin and 52 were profiled from a non-primary specimen (lymph node 25, other 11, lung 10, liver 5) [these sum to 51; one count requires correction]. The remaining 288 patients constitute the primary-site analytic cohort, comprising 195 bladder-primary and 93 upper tract primary tumors. Counts of program-low and program-high tumors and of response-evaluable patients are shown for each site.

**(b)** Distribution of tsMHC-II program GSVA scores across the 288-sample analytic cohort, with the fitted two-component Gaussian mixture (component densities in red, summed density in green) and the cutpoint defined by the intersection of the component densities (0.007, vertical line). Scores were recomputed on the 288-sample matrix rather than subset from a larger run, because GSVA scores each sample relative to the gene distributions of the input matrix.

**(c)** Overall survival in the full 288-patient cohort without site restriction, stratified by program status. Log-rank p = 0.28; hazard ratio 0.85 (95% CI 0.64 to 1.14, p = 0.277). No association is detectable when tumors of both primary sites are pooled.

**(d)** Forest plot of the bladder-primary overall survival model ladder. Each row shows the hazard ratio for program-high versus program-low with 95% confidence interval, together with the number of patients and events. Rows comprise the pooled both-site model, the unadjusted bladder model, the unadjusted model restricted to patients with complete clinical covariate data, the model adjusted for ECOG performance status, liver metastasis, prior platinum therapy, and sex, the unadjusted model restricted to patients with evaluable tumor mutational burden, the model adjusted for mutational burden, and the model adjusted for mutational burden and PD-L1 immune-cell staining. Comparison of rows 3 and 4, which are fitted on the same 177 patients, shows that the attenuation after clinical adjustment reflects confounding rather than reduced sample size. Program-low is the reference category throughout.

**(e)** Site-specific hazard ratios across three model specifications, on the same axis convention as panel d, with the site-by-program interaction p value shown for each specification. The interaction is significant in the unadjusted cohort (p = 0.0018) and in complete cases (p = 0.021), and attenuates after adjustment for clinical prognostic factors (p = 0.059). The interaction term is a ratio of hazard ratios and is not directionally comparable to the within-site estimates.

**(f)** Objective response by tsMHC-II program status and primary site among response-evaluable patients (bladder n = 168; upper tract n = 80). Bars show the proportion achieving complete or partial response versus stable or progressive disease, with counts and percentages overlaid. Bladder p = 0.154; upper tract p = 0.018.

#### SUPPLEMENTARY TABLES

**Supplementary Table 1.** Bladder cancer cell lines profiled in this study, listing the ATCC designation, molecular subtype, the assays performed for each line (RNA-seq, ATAC-seq, and surface MHC-II flow cytometry), and notes on histology and Cellosaurus identifier.

**Supplementary Table 2.** The consensus in vitro IFN-γ response signature, giving the genes significantly upregulated (UP) and downregulated (DOWN) after IFN-γ stimulation across the responsive bladder cancer cell lines. This signature is tested for enrichment in patient tumor cells in Figure 2e.

**Supplementary Table 3.** The in vitro arm of the tsMHC-II program signature: genes significantly upregulated by IFN-γ stimulation in at least five of six bladder cancer cell lines, used as the first of three inputs to the signature intersection.

**Supplementary Table 4.** Genes significantly upregulated in tsMHC-II-positive versus tsMHC-II-negative malignant epithelial cells in the internal NMIBC snRNA-seq cohort, used as the second input to the signature intersection.

**Supplementary Table 5.** Genes significantly upregulated in tsMHC-II-positive versus tsMHC-II-negative malignant epithelial cells in the external MIBC snRNA-seq cohort, used as the third input to the signature intersection.

**Supplementary Table 6.** The final 11-gene tsMHC-II program signature, defined as the three-way intersection of the gene sets in Supplementary Tables 3 to 5: CD74, CIITA, CXCL10, CXCL11, ICAM1, GBP5, IL32, SOD2, HAPLN3, TNFRSF1B, and SLC12A7.

**Supplementary Table 7.** Clinical, molecular, and immune characteristics of the 195 bladder-primary patients in the IMvigor210 analytic cohort, compared between program-low (n = 106) and program-high (n = 89) tumors. The four covariates used in the adjusted survival model (ECOG performance status, liver metastasis, prior platinum therapy, and sex) are closely balanced between groups. Two variables are not: metastatic disease status, driven by a higher frequency of lymph-node-only disease in program-high tumors, and tumor mutational burden.

**Supplementary Table 8.** Clinical, molecular, and immune characteristics for all 288 patients in the analytic cohort, compared between bladder-primary (n = 195) and upper tract primary (n = 93) tumors. The two groups are comparable in sex, performance status, tumor mutational burden, PD-L1 immune-cell level, and both Lund and TCGA subtype composition, and differ materially only in prior platinum exposure.
